# *Fusobacterium nucleatum* is enriched in invasive biofilms in colorectal cancer

**DOI:** 10.1101/2024.12.30.630810

**Authors:** Jessica Queen, Zam Cing, Hana Minsky, Asmita Nandi, Taylor Southward, Jacqueline Ferri, Madison McMann, Thevambiga Iyadorai, Jamuna Vadivelu, April Roslani, Mun Fai Loke, Jane Wanyiri, James R. White, Julia L. Drewes, Cynthia L. Sears

## Abstract

*Fusobacterium nucleatum* is an oral bacterium known to colonize colorectal tumors, where it is thought to play an important role in cancer progression. Recent advances in sequencing and phenotyping of *F. nucleatum* have revealed important differences at the subspecies level, but whether these differences impact the overall tumor ecology, and tumorigenesis itself, remain poorly understood. In this study, we sought to characterize *Fusobacteria* in the tumor microbiome of a cohort of individuals with CRC through a combination of molecular, spatial, and microbiologic analyses. We assessed for relative abundance of *F. nucleatum* in tumors compared to paired normal tissue, and correlated abundance with clinical and pathological features. We demonstrate striking enrichment of *F. nucleatum* and the recently discovered subspecies *animalis* clade 2 (Fna C2) specifically in colon tumors that have biofilms, highlighting the importance of complex community partnerships in the pathogenesis of this important organism.

## Introduction

Colorectal cancer (CRC) is a leading cause of cancer morbidity and mortality, with 1.8 million new cases and >800,000 deaths annually ^1^. Microbial dysbiosis is increasingly recognized as a factor in colon tumor initiation and progression. This dysbiosis can manifest as enrichment or depletion of specific bacterial taxa or in alterations of bacterial communities, such as the development of mucus-invasive biofilms ^2^. *Fusobacterium nucleatum*, a Gram-negative anaerobe and common member of the human oral microbiota, is known to be associated with CRC ^3–6^. Clinical studies from North American, European, and Asian cohorts have demonstrated enrichment of *F. nucleatum* in a subset of CRC compared to healthy colon tissues ^3–6^. Data from animal models and translational studies have suggested potential roles for this organism in colon tumor development, progression, metastasis and/or treatment response ^7^. In the mouth, *F. nucleatum* plays an important role as a bridging species linking early and late colonizers of polymicrobial biofilms^8^. Biofilms have also been recognized as common features of colon tumors ^9,10^, where it is posited that *F. nucleatum* and other organisms invade into the mucosa to contact host cells and promote inflammation and tumorigenesis. *F. nucleatum* is a heterogenous species consisting of four subspecies anticipated to become distinct species: *animalis*, *nucleatum*, *polymorphum*, and *vincentii* ^11^. Recently, subspecies *animalis* has been further classified into two clades, with identification of clade Fna C2 as prominently featured in the colon tumor microenvironment ^12^. In this study, we sought to characterize the tumor microbiome of a large cohort of individuals with CRC through a combination of sequencing and spatial analysis, and to identify features associated with enrichment of *F. nucleatum* and its subspecies and clades.

## Results

### Colon tumors frequently have polymicrobial biofilms containing Fusobacteria

The study population consisted of 116 individuals diagnosed with colorectal cancer in Malaysia, from whom paired tumor and distant normal colon tissues were collected at the time of CRC tumor resection surgery (**Fig 1A**). Four of these individuals underwent surgery for resection of masses later identified as adenomas (polyps). This 116 person cohort is inclusive of two previously published smaller cohorts (MAL1 and MAL2) ^10^, to which we added an additional 72 individuals with CRC, yielding an assessment of 143 additional tumor and/or paired normal samples (MAL3). Demographic and clinical data collected included age, sex, ethnicity, tumor location, and tumor stage (**Fig 1B**). The proximal colon through the hepatic flexure was defined as right colon, and distal to the hepatic flexure as left colon, as previously described ^10^. To assess for the presence of biofilms, we performed fluorescence in situ hybridization (FISH) on methacarn-fixed tissues of colon tumor and paired normal samples (**Fig 2A**). We found that biofilms were prevalent on colon tumors, visualized in 68.8% (77/111) (**Fig 2B**). There was high concordance in biofilm status between tumors and paired normal tissue; in individuals who had biofilms observed on their tumors, 79.0% (60/76) also had biofilms on distant normal tissue (**Fig 2C**). In those with no biofilms visualized on their tumors (n=34), there were no biofilms observed in paired normal tissue. Biofilms were more frequently observed on right-sided tumors (33/36; 91.7%) compared to left (44/75; 58.7%; p=0.0003), with no significant association with tumor stage (**Fig 2D-E**). Of 70 biofilm-positive tumors available for multi-probe FISH, we identified broad patterns of microbial biofilm community membership, consisting of either polymicrobial biofilms (94.3%; 66/70) with or without *Fusobacterium* (genus), or, rarely, biofilms predominated by proteobacteria (5.7%; 4/70) (**Fig 2F-G**). Among the 66 polymicrobial biofilms, we visualized *Fusobacterium spp.* in 68.2% (45/65), ranging from sparse populations of individual rods to dense blooms where *Fusobacterium* was the predominant organism. Although biofilms were not associated with tumor stage (**Fig 2E**), the presence of *Fusobacterium* in tumor biofilms was associated with later tumor stage compared to polymicrobial biofilms where *Fusobacterium* was absent (p=0.0212) (**Fig 2F**).

**Figure 1.**
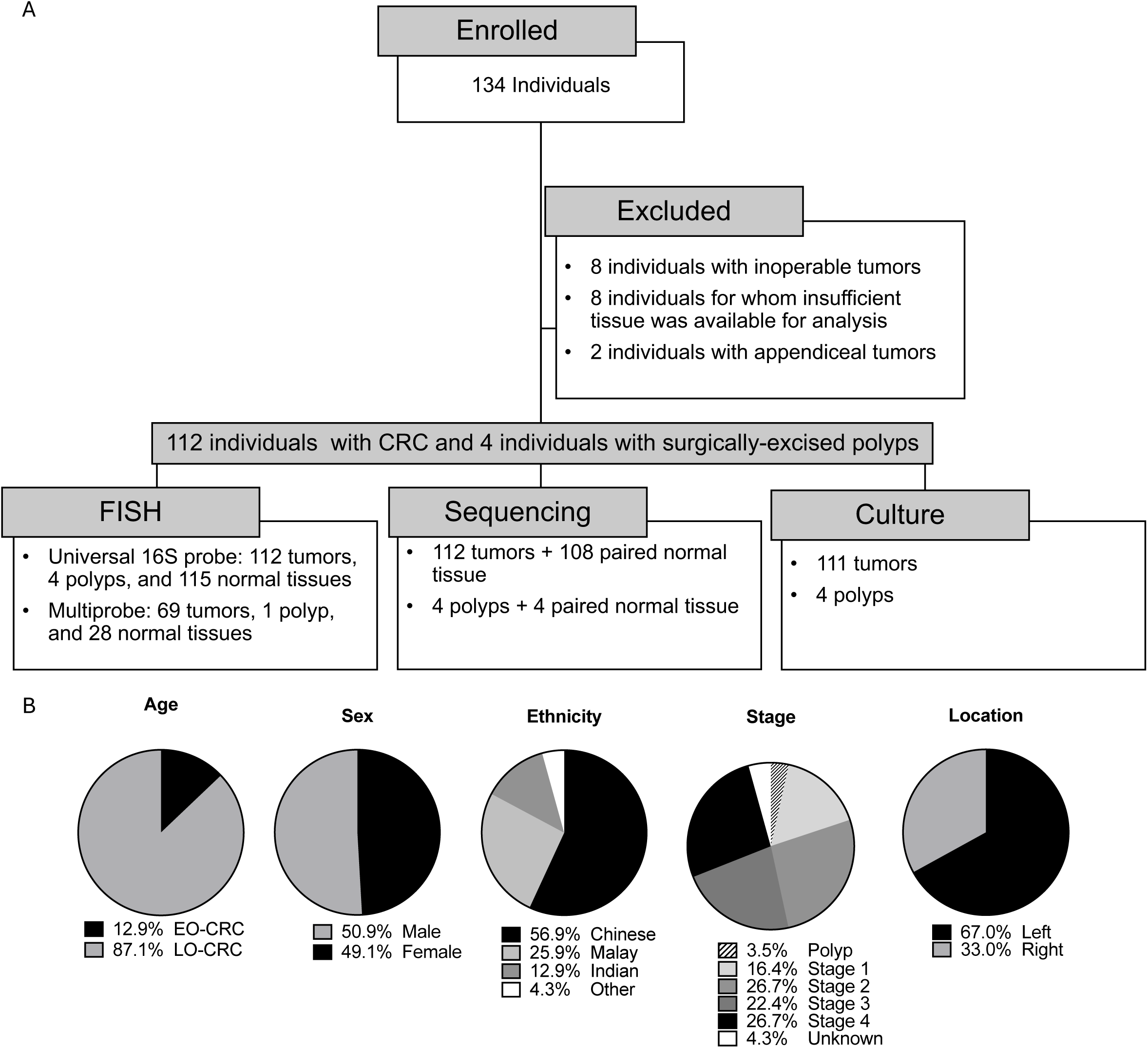
The CRC cohort. A) Flow diagram of study participants and downstream analysis of samples. B) Demographic characteristics of the cohort (n=116), including age, sex, ethnicity, stage, and tumor location. EO-CRC: early-onset colorectal cancer, defined as age under 50. LO-CRC: late-onset colorectal cancer, defined as age 50 or above.

**Figure 2.**
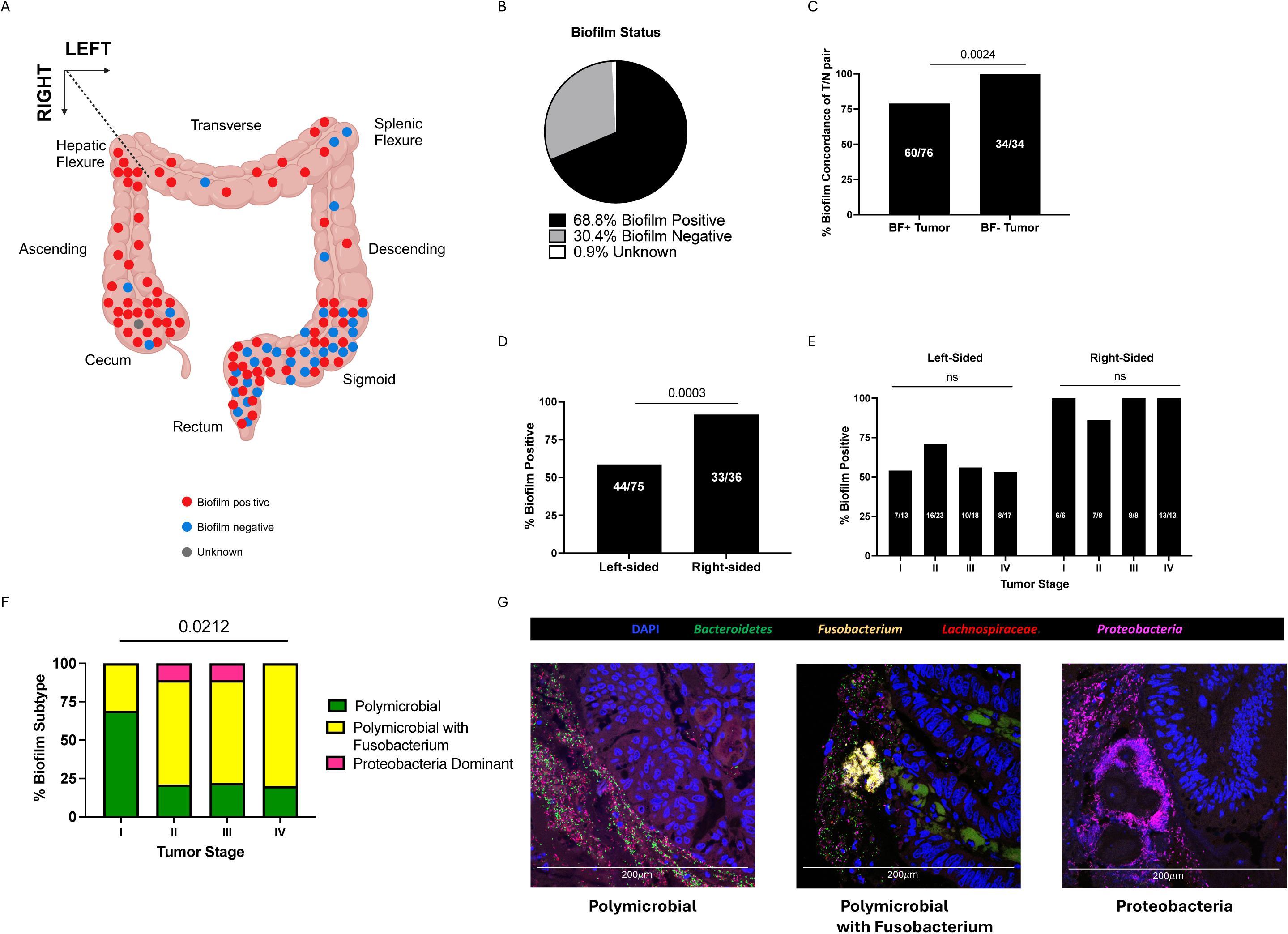
Biofilm analysis of the CRC cohort. A) The location and biofilm status of CRC tumors overlaid on a diagram of a colon. One tumor marked as unknown had no fixed tissue suitable for screening. B) Biofilm status of CRC tumors classified as either positive or negative by FISH. C) Percent of left and right-sided tumors positive for biofilms. D) Percent concordance of biofilm status in paired tumor (T) and normal (N) tissue for biofilm-positive (BF+) tumors and biofilm-negative (BF-) tumors. E) Percent of tumors positive for biofilms stratified by tumor location and stage. C-E analyzed by Fisher’s exact test, with p<0.05 considered significant. f) Stacked bar graph depicting percent of tumor biofilms of each subtype screened by multi-probe FISH (n=70) and stratified by tumor stage. Analyzed by nonparametric Kruskal-Wallis one-way ANOVA, with p<0.05 considered significant. G) Representative 40X images of multi-probe FISH of each biofilm subtype stained for DAPI (blue), Bacteroidetes (green), Fusobacterium (yellow), Lachnospiraceae (red), and Proteobacteria (pink).

### *Fusobacteria* are enriched in the colon tumor microbiome

We performed 16S rRNA amplicon sequencing on tumor biopsies and paired normal tissue to identify bacterial taxa enriched or depleted in the tumor microenvironment. Two subsets of this cohort were previously analyzed by 16S rRNA amplicon sequencing and published ^10^: MAL1 (consisting of 22 tumors and 1 adenoma with 20 paired normal samples) and MAL2 (consisting of 24 tumors with 22 paired normal samples, 3 of which were replicate pairs from MAL1). We performed 16S rRNA amplicon sequencing on an additional 79 tumors, 3 adenomas, and 71 paired normal samples; this subset, termed MAL3, included 10 replicate tumors previously sequenced in the MAL1 and/or 2 cohorts. Although all 3 cohorts were sequenced independently, each was sequenced using the V3-V4 hypervariable region as the amplification target, and data were analyzed using the same computational pipeline to assign species-level taxonomic identification (Resphera Insight, Baltimore, MD). Analysis of these 10 replicates demonstrated consistent taxonomic assignments across independent sequencing runs for replicate samples at the phylum through species levels (**Table S1A-G; Fig S1A-E; Fig S2**), including *Fusobacterium* species (**Fig S3**), which are the particular focus of this study. Given the concordance of taxonomic assignments between replicates in differing sequencing runs, we pooled MAL1, MAL2, and MAL3 data for all subsequent analyses (see **Table S2** for all taxonomic assignments). Altogether, the combined MAL cohort presented herein consists of 108 unique tumor and normal tissue pairs, 4 adenoma and normal tissue pairs, and an additional 4 tumors for which paired normal sequencing data is unavailable.

Consistent with prior studies ^13,14^, tumors had lower alpha diversity than normal tissue as measured by the number of observed species (p = 0.0042). Biofilm-negative tumors had lower alpha diversity than biofilm-negative normal tissue, as measured by number of observed species (p = 0.019) or Chao Index (p = 0.0346) (**Fig S4A**). Although biofilm-positive tumors were more diverse than biofilm-negative tumors by Chao Index (p = 0.0096), there was no significant difference in alpha diversity between biofilm-positive tumor and normal samples, by Chao Index, Shannon Index, Simpson Index, or number of observed species. Neither sample type (tumor vs normal) nor biofilm status (positive vs negative) was associated with differences in beta diversity, as measured by Bray-Curtis dissimilarity, with significance estimated by Permutational multivariate analysis of variance (PERMANOVA) (**Fig S4B**). *F. nucleatum* was present in 86.6% of CRC tumors in this cohort (97/112), with relative abundance ranging from 0.01% to 40% of total microbial reads (**Table S2**). Analysis of the 108 tumor/normal pairs revealed significant enrichment of *F. nucleatum* in the tumor microbiome compared to paired normal samples (p<0.0001) (**Fig 3**). Resphera Insight attempts to achieve species-level resolution; however, when a confident single species assignment is not feasible, the method minimizes false positives by providing ambiguous assignments that may include multiple closely related species. This significant enrichment in the tumor microenvironment was observed whether *F. nucleatum* abundance was calculated using confident (**Fig 3**) or confident + ambiguous (**Fig S5A**) assignments. Other *Fusobacterium* species enriched in tumors included *F. peridonticum* (p=0.0005), *F. simiae* (p=0.0293), and *F. varium* (p=0.0094) (**Fig 3**). The latter has previously been found to be enriched only in Southern Chinese populations ^15^. In our cohort, *F. varium* was enriched in tumors vs paired normal tissue only for individuals of Malay descent (**Fig S5B**). Also of note, *Bacteroides fragilis* was significantly enriched in the tumor microbiome relative to paired normal tissue (p=0.0002), whereas other taxa previously linked to the CRC microbiome were not enriched in our cohort, including *Akkermansia muciniphila, Enterococcus faecalis*, and *Streptococcus gallolyticus* (**Fig S6**). We also observed relative depletion of *Alistipes onderdonkii* (p=0.0261), *Bifidobacterium longum* (p=0.0029), *Parabacteroides distasonis* (p=0.0149), and *Ruminococcus bromii* (p<0.0001), with a trend toward depletion of *Alistipes senegalensis* (p=0.0620) in tumor samples.

**Figure 3.**
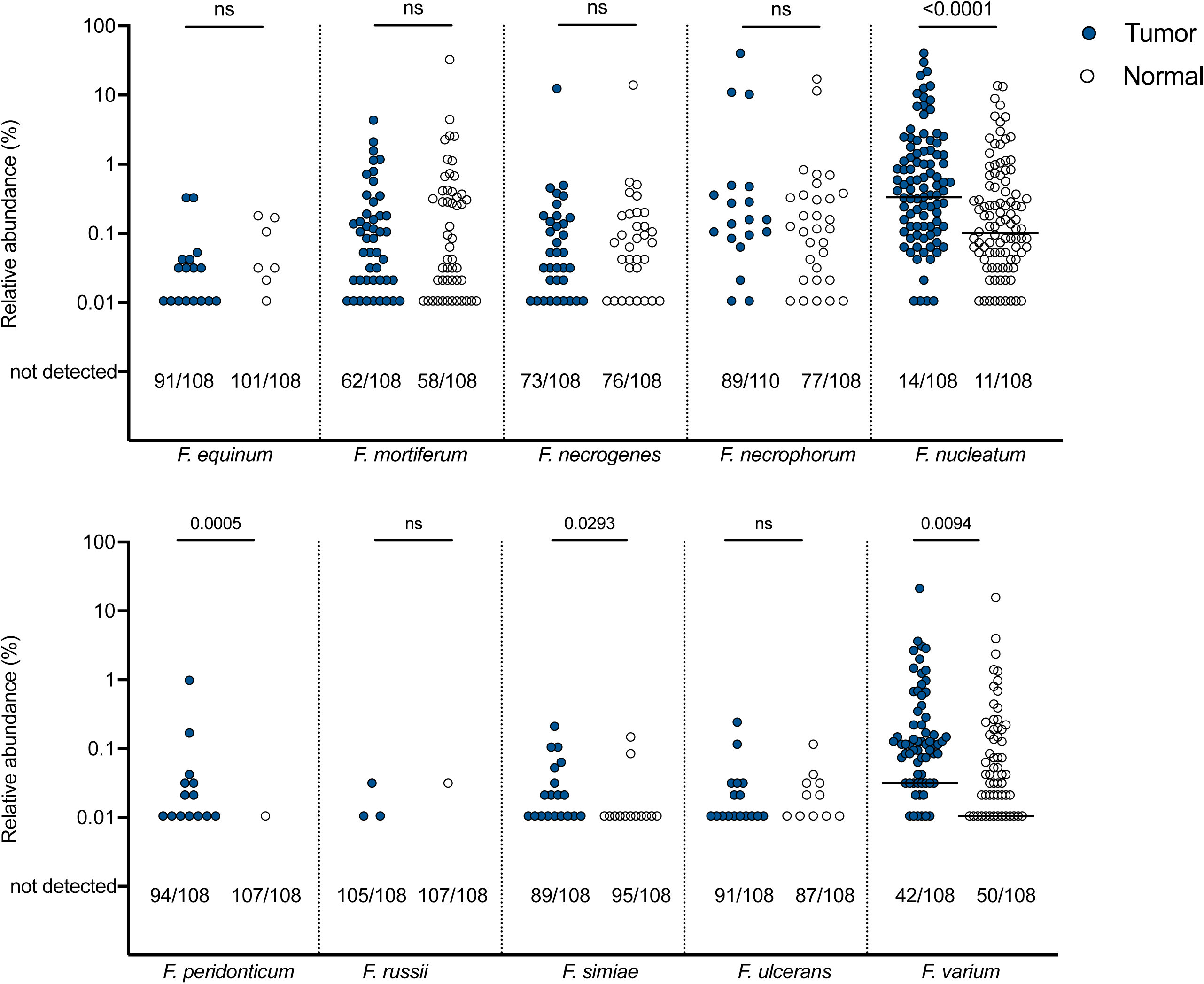
Abundance of *Fusobacterium* species in tumor/normal pairs. Relative abundance on a log scale by 16S rRNA amplicon sequencing of each depicted *Fusobacterium* species in pairs of tumor (blue circle) and normal (clear circle) tissues. Bars indicated median. Numbers below each graph depict the number of samples out of the total in which each species was not detected by sequencing (i.e. 0% abundance). Analyzed by two-tailed Wilcoxon matched-pairs signed rank test, with p<0.05 considered significant. See Fig S5 for abundance of *F. nucleatum* as a sum of confident and ambiguous species assignments, and for *F. varium* abundance stratified by ethnicity.

### *Fusobacterium nucleatum* and other oral organisms are associated with colon tumor biofilms

We next investigated individual and tumor characteristics associated with *F. nucleatum* abundance in the tumor microbiome. Enrichment of *F. nucleatum* in this cohort was independent of sex or tumor location, but was seen among late-onset (LO) CRC and Chinese and Malay individuals. In contrast, no enrichment was seen among individuals with early-onset (EO) CRC or those of Indian descent (**Fig 4A-D**). Notably, *F. nucleatum* enrichment was observed only in later cancer stages (p=0.0041, 0.1037, and 0.0046, respectively, for stages II-IV) (**Fig 4E**) and in biofilm-positive tumors (p<0.0001) (**Fig 4F; Fig S5C**. This enrichment in later cancer stages is in fact driven by higher abundance of *F. nucleatum* in tumors that have biofilms (p=0.0012, 0.0084, and p<0.0001, respectively, for biofilm-positive tumors at stages II-IV, whereas biofilm-negative tumors had no significant enrichment at any cancer stage (**Fig 4G**). This finding supports our observations from multiprobe FISH (**Fig 2F**), highlighting that although *F. nucleatum* is present in a majority of tumors in this cohort, it is specifically enriched in association with biofilms in advanced disease. Because of the association of *F. nucleatum* with oral biofilms, we assessed for enrichment of other oral microbiome community members. 16S rRNA amplicon sequencing analysis revealed enrichment of *Actinomyces odontolyticus* (p=0.0127), *Gemella morbillorum* (p<0.0001)*, Haemophilus influenzae* (p=0.0164), *Lachnoanaerobaculum orale* (p<0.0001), *Parvimonas micra* (p *=*0.0255), and *Porphyromonas gingivalis* (p=0.0057) in the tumor microbiome (**Fig S7**). Of these organisms, the relative abundance of *G. morbillorum* (p=0.005), *L. orale* (p=0.006), *P. micra* (p<0.0001), and *P. stomatis* (p<0.0001) were positively associated with abundance of *F. nucleatum* in the tumor microbiome (**Fig 4H**).

**Figure 4.**
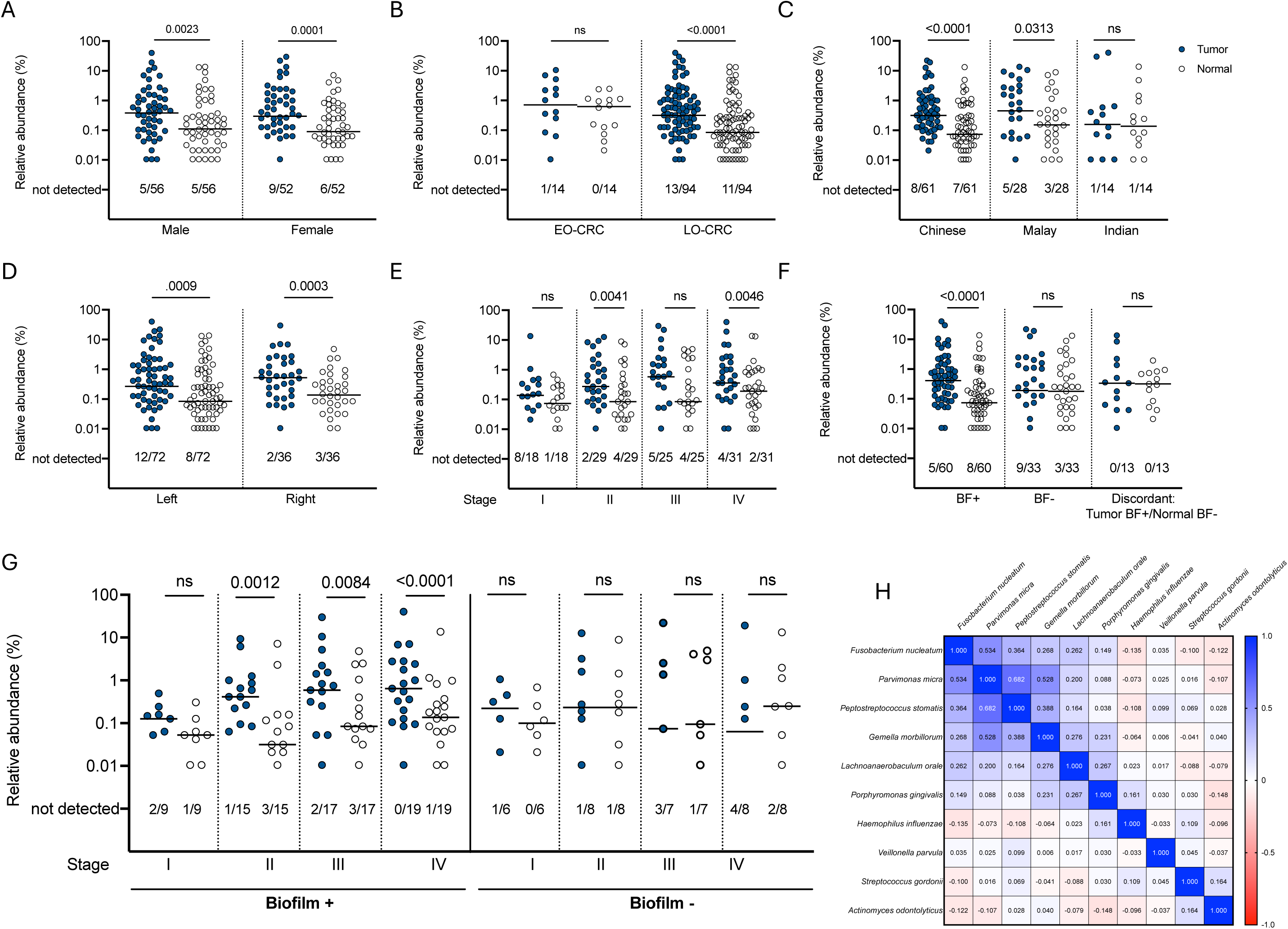
*Fusobacterium nucleatum* abundance by individual and tumor characteristics. Relative abundance by 16S rRNA amplicon sequencing of *Fusobacterium nucleatum* in pairs of tumor (blue circle) and normal (clear circle), stratified by A) sex, B) age, C) ethnicity, D) tumor location, E) tumor stage, F) biofilm status, and G) both biofilm status and tumor stage. Bars indicated median. Numbers below each graph depict the number of samples out of the total in which *F. nucleatum* was not detected by sequencing. Analyzed by two-tailed Wilcoxon matched-pairs signed rank test, with p<0.05 considered significant. EO-CRC: early-onset colorectal cancer. LO-CRC: late-onset colorectal cancer. See Fig S5 for abundance of *F. nucleatum* as a sum of confident and ambiguous species assignments, stratified by biofilm status. H) Matrix of *F. nucleatum* and other oral biofilm organism abundances in tumor samples, analyzed by non-parametric Spearman correlation coefficient. See Fig S7 for relative abundance of each organism in the matrix in tumor/normal pairs.

### *F. nucleatum subspecies animalis* clade 2 is the predominant *Fusobacterium* enriched in colon tumors with biofilms

We next assessed the full cohort for the presence of the recently delineated clades within subspecies *animalis* using clade-specific amplicon sequence variants, as previously described ^12^. We found that both subspecies *animalis* clade 1 (Fna C1) and clade 2 (Fna C2) were significantly enriched in the tumor microenvironment compared to normal tissues (p=0.0046 and p<0.0001, respectively) (**Fig 5A**). However, Fna C2 was strikingly enriched in both tumors and normal tissues compared to Fna C1 (p<0.0001). Similar to our findings at the species level (**Fig 4F**), when compared to paired normal tissues, Fna C1 and C2 are only enriched in tumors that have biofilms (p=0.0027 and p<0.0001, respectively) (**Fig 5B**). When biofilm-positive tumors were stratified by stage, Fna C1 only reached statistically significant enrichment in stage 3 tumors (**Fig 5C**), whereas Fna C2 was significantly enriched at all tumor stages (**Fig 5D**). In our small cohort of 4 individuals with surgical adenomatous polyps, 2 had biofilms and 2 did not (**Fig S8A**). *F. nucleatum*, including Fna C1 and Fna C2, were detected in a majority of these adenomas (**Fig S8B**), although the small sample size precluded comparative analysis. Altogether, we find striking enrichment of Fna C2 in tumors that span the adenoma to advanced carcinoma sequence, and particularly in tumors displaying biofilms.

**Figure 5.**
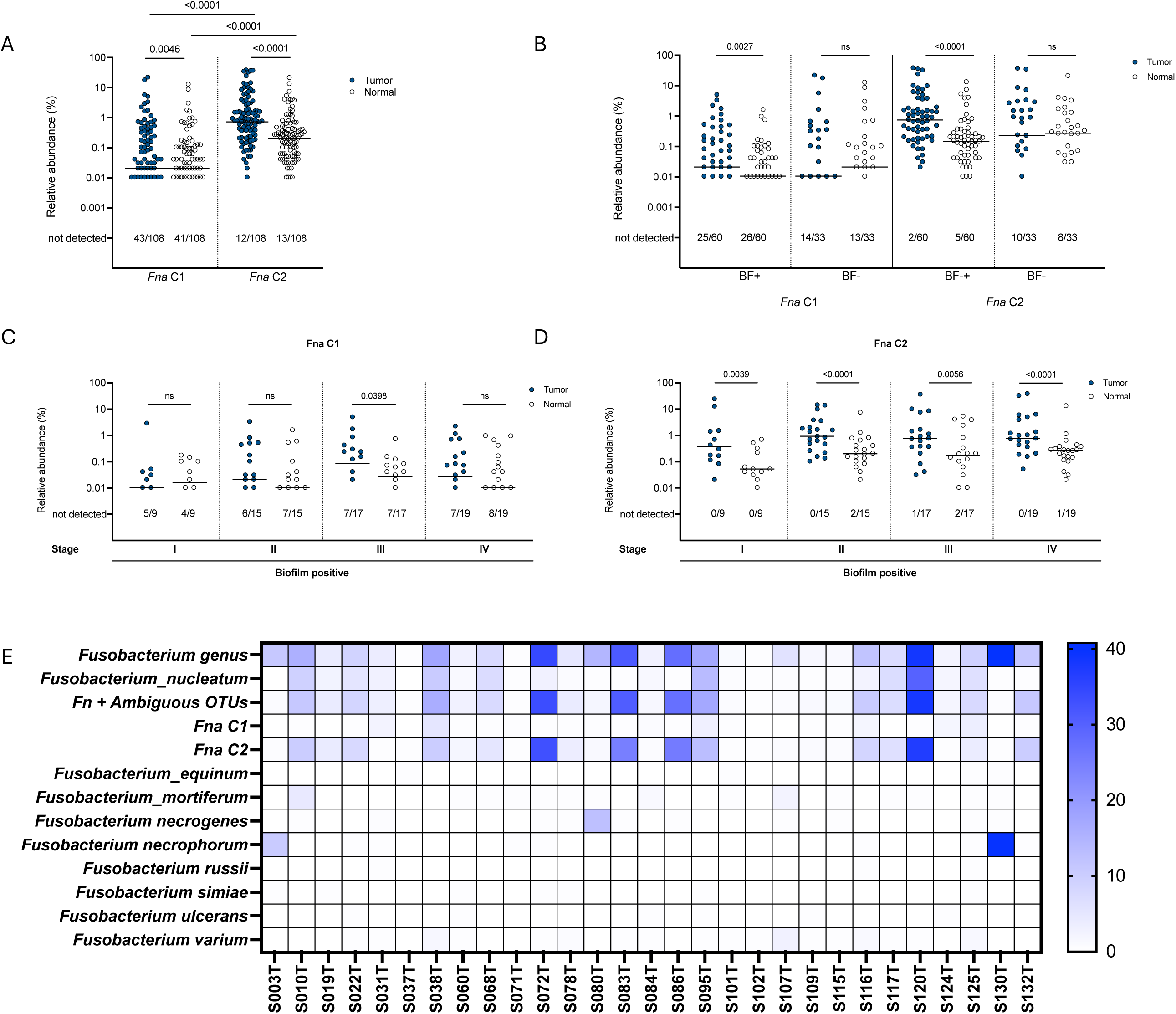
*Fusobacterium nucleatum subspecies animalis* abundance by clade and correlated with FISH. A) Relative abundance by 16S rRNA amplicon sequencing of *Fusobacterium nucleatum subspecies animalis* clade 1 (Fna C1) and clade 2 (Fna C2) in pairs of tumor (blue circle) and normal (clear circle). Tumor/normal pairs analyzed by two-tailed Wilcoxon matched-pairs signed rank test, and tumor/tumor or normal/normal comparisons analyzed by two-tailed Mann Whitney test, with p<0.05 considered significant. B) Relative abundance of Fna C1 and C2 stratified by biofilm (BF) status and C-D) by both biofilm status and tumor stage. Bars indicated median. Numbers below each graph depict the number of samples out of the total in which each clade was not detected by sequencing. E) Heatmap depicting relative abundance by 16S rRNA amplicon sequencing for the *Fusobacterium* genus and each labeled *Fusobacterium* species or clade for n=29 tumors where dense blooms of Fusobacteria were visualized by FISH.

We assessed whether Fna C2 abundance by 16S rRNA amplicon sequencing correlated with the density of *Fusobacterium* visualized by FISH (**Fig 5E**). Of note, the *Fusobacterium* FISH probe is genus-specific. In tumor biofilms with dense blooms of Fusobacteria (n=29), a small number of tumors had higher abundance of non*-nucleatum* Fusobacteria by sequencing analysis. However, in general, the abundance of Fna C2 was similar to the total estimated abundance of *F. nucleatum* (confident + ambiguous assignments) (**Table S2**), consistent with Fna C2 being dominant in the tumor microenvironment.

To identify whether other *F. nucleatum* subspecies were detectable in CRC tissues and to cross compare methodologies, we complemented our 16S rRNA amplicon sequencing analysis with a PCR-based method for distinguishing the *F. nucleatum* subspecies ^16^. Of 34 tumors tested, *F. nucleatum* was detectable by PCR in 25, and among these, subspecies *animalis* was most frequently detected (**Fig S9A**), consistent with previous reports ^16,17^. When stratified by biofilm status, subspecies *animalis* was most frequently detected in biofilm positive tumors, whereas subspecies *vincentii* was more frequently detected in biofilm negative tumors (**Fig S9B**). Notably, multiple subspecies were detected in 20% of tumor biopsies tested (5/25).

In parallel to sequencing and PCR analysis of *Fusobacterium* in the tumor microbiome of our cohort, we also used selective culture media to isolate unique *Fusobacterium* strains. Of the 13 *Fusobacterium* strains isolated, 6 were *F. nucleatum,* 50% of which were subspecies *animalis* (**Table S3**).

### Fna C2 abundance correlates with predicted metabolic shifts

To begin to investigate functional consequences of Fna C2 enrichment and biofilm formation in the tumor microenvironment, we used the PICRUSt2 computational analysis pipeline to predict differences in metabolic pathways inferred from our 16S rRNA amplicon data (**Fig 6; Fig S10; Fig S11; Table S2I**). We identified 83 pathways that predicted differential metabolic function between tumor samples with low Fna C2 abundance (<1%) and high abundance (≥1%) (**Fig 6**). Many of these pathways are involved in amino acid and cofactor biosynthesis, carbohydrate degradation, energy metabolism, or represent metabolic superpathways. Fna C2 abundance was positively associated with a small number of PICRUSt2 pathways (**Fig S10**), notably including L-1,2-propanediol degradation (PWY-7013), consistent with the previous observation that Fna C2 has conservation of genetic content associated with 1,2-propanediol (1,2-PD) metabolism ^12^. We also identified positive association of Fna C2 abundance with amino acid metabolism, including L-glutamate degradation V (P162-PWY) and L-lysine fermentation to acetate and butanoate (P163-PWY). Fna C2 abundance was also positively associated with protein N-glycosylation, which is known to be aberrant in tumors and is potentially a critical factor in tumor immune evasion ^18^. A large number of inferred metabolic functions were negatively associated with Fna C2 abundance (**Fig S10**). These included numerous carbohydrate metabolism pathways, such as the superpathway of fucose and rhamnose degradation (FUC_RHAMCAT-PWY) and other related pathways (FUCCAT-PWY, RHAMCAT-PWY), and sucrose degradation IV (PWY-5384).

**Figure 6.**
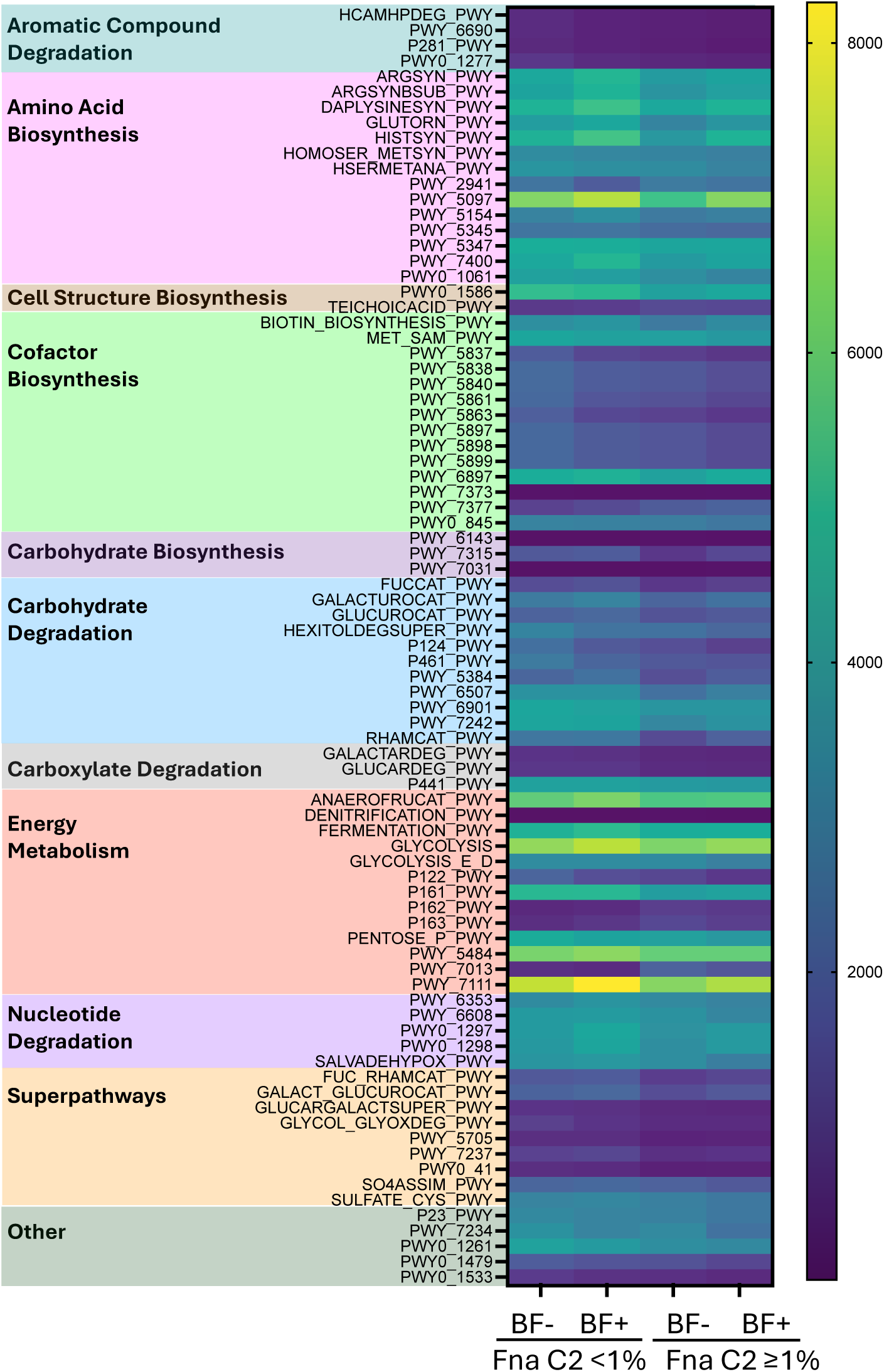
PICRUSt2 pathways associated with Fna C2 abundance in CRC tumors. Heatmap of all statistically significant PICRUSt2 pathways with differences in gene content between low Fna C2 abundance (<1%) and high abundance (≥1%) in tumors (n=112), as analyzed by Mann Whitney test, with p<0.05 considered significant. Data are further stratified by biofilm (BF) status within each abundance category.

Lastly, we sought to investigate whether Fna C1 abundance was associated with a metabolic signature similar to Fna C2. When Fna C1 is present in the tumor microenvironment, it is typically at low abundance (median 0.02%) and absent in nearly 40% of tumors (**Fig 5A**). We therefore sought to identify PICRUSt2 pathways significantly associated with presence or absence of Fna C1. We identified 34 pathways differentially represented in tumors with or without Fna C1, with many pathways associated with aromatic compound degradation and carbohydrate metabolism (**Fig S11**). Of these 34 pathways, only 5 pathways were significantly associated with both Fna C1 and Fna C2 abundance: L-glutamate degradation V (P162-PWY), L-1,2-propanediol degradation (PWY-7013), protein N-glycosylation (PWY-7031), tRNA processing (PWY0-1479), and teichoic acid biosynthesis (TEICHOCACID-PWY).

## Discussion

*Fusobacterium nucleatum*, which typically resides in the oropharynx, has been consistently linked to the colon cancer microbiome in numerous studies across diverse cohorts ^3,4,19–22^. *F. nucleatum* enrichment has been associated with specific pathological and clinical features such as high levels of microsatellite instability (MSI-high), the CpG island methylator phenotype (CIMP), and poor prognosis ^23–26^. Controversy remains as to the precise role(s) *F. nucleatum* plays in tumor initiation, progression, and immune evasion, although a large body of data argues against it merely existing as a passive colonizer of colon tumors. One key barrier to uncovering critical host-bacterial interactions in the tumor microenvironment is the diversity of *F. nucleatum* strains employed to date in mechanistic studies that may not accurately reflect the genomic content or adhesion factor repertoire of CRC-associated strains. Therefore, a key question has been which subspecies of this diverse organism are enriched in colon tumors. Inconsistent phenotypes in mouse models have also raised the question as to the critical community partners with which *F. nucleatum* may collaborate to colonize the lower gastrointestinal tract and promote carcinogenesis ^27–29^.

In this large cohort of individuals with CRC, we report that *F. nucleatum* is significantly enriched in the colon tumor microbiome when compared to distant normal tissues. A combination of sequencing, culture, and spatial evaluation has, in particular, allowed us to link *F. nucleatum* abundance in the tumor microenvironment with the presence of tumor biofilms as well as with enrichment of other oral microbiome members, particularly in advanced cancer stages. These data build on previous analyses linking *F. nucleatum* to CRC biofilms ^10^, and confirm the finding that biofilms on colon tumors frequently contain *Fusobacteria*. Although *F. nucleatum* has previously been linked to CRC biofilms, we report the surprising finding that *F. nucleatum* is enriched *only* in colon tumors that have biofilms. By combining microbial sequencing and spatial analysis, we find that *F. nucleatum*, dominated by Fna C2, is frequently present in CRC biofilms and can bloom into dense populations, the latter rarely identified in biofilms on normal tissue ^10^. This observation supports the theory that *F. nucleatum* is itself not a tumor initiator, but blooms in the tumor environment under specific conditions that allow it to flourish and subsequently promote tumor progression. PICRUSt2 analysis suggests that Fna C2 abundance in the tumor microenvironment may be associated with altered metabolic function, a key hypothesis for future study. Consistent with previous studies, we also find enrichment of other oral organisms in colon tumors, suggesting that *F. nucleatum* expansion occurs in the context of an oral microbiome community ^10,30,31^.

We find that subspecies *animalis* is most frequently detected by PCR and isolated by culture, consistent with other reports ^16,17^. One limitation of our study was inability to perform strain-level comparisons of *Fusobacterium* isolates because of low culture yield, generating a library of only 13 Fusobacteria strains from the cohort of 112 individuals. Culture yield was likely impacted by long-term storage of tumor biopsies for 5-10 years prior to attempts at isolating *Fusobacterium* strains. Recent work has identified two clades (Fna C1 and C2) within subspecies *animalis* that were frequently detected in the mouths of healthy individuals, whereas only clade Fna C2 was enriched in colon tumors ^12^. We confirm this finding in our cohort, with Fna C2 representing the most abundant *Fusobacterium* in the CRC microbiome, although our data show that other *Fusobacterium* species are also significantly enriched in the tumor microenvironment. Although we consider Fna C2 a high priority group for future investigation, fusobacterial communities in CRCs are complex, strongly suggesting that investigation of community features beyond Fna C2 is required to understand the mechanistic determinants of biofilms, *Fusobacterium,* and CRC.

Notably, among the subset of tumors with *Fusobacterium* blooms, the overall abundance of *F. nucleatum* and other *Fusobacterium* species ranged in the tumor microenvironment from 0.06 to 40.8% of total microbial reads, suggesting that there are concentrations of Fusobacteria in specific niches in the tumor microenvironment that are underappreciated by non-spatial sequencing techniques. Future investigations into the role of *F. nucleatum* in CRC pathogenesis will need to harness emerging spatial omics analyses to better understand the specific interactions of *F. nucleatum* and biofilm community members with host cells.

These analyses build on existing data demonstrating an association of *F. nucleatum* with the colon tumor microenvironment, and add critical resolution that highlights the abundance of *F. nucleatum*, and specifically subspecies *animalis* clade 2, in mucus-invasive biofilms observed particularly in late-stage disease. Our findings suggest that *F. nucleatum*:bacterial interactions within mucus-invasive biofilms are understudied features of CRC carcinogenesis.

## Acknowledgments

We honor the memory of the late Dr. Khean Lee Goh, who contributed to this work. We thank Kumar Thulasi, Han Ming Gan, Hoong Yin Chong, and Sandip Kumar for their technical assistance.

We thank the Sidney Kimmel Comprehensive Cancer Center and the Johns Hopkins University Oncology Tissue Services (supported by NCI grant P30 CA006973), and the Johns Hopkins University Institute for Basic Biomedical Science Microscopy Facility for use of their Zeiss LSM 780 Confocal microscope (with Fluorescence Correlation Spectroscopy) (supported by NIH Grant S10 OD016374).

This work was supported by the National Institute of Allergy and Infectious Diseases training grant T32-A1007291, the Biocodex Microbiota Foundation, the Burroughs Wellcome Fund Career Award for Medical Scientists [1022128], and the Black in Cancer award sponsored by the Emerald Foundation, Inc. (J.Q.); The U-RISE Program at the University of Maryland, Baltimore County (UMBC), which is supported by the National Institute of General Medical Sciences of the National Institutes of Health under Award Number T34-GM136497 (Z.C.); The National Institute of General Medical Sciences training grant 3-T32-GM136577 (H.M.); the University of Malaya Research Grant RP016A-13HTM (J.V.); National Cancer Institute grant R00 CA230192 (J.L.D.); the Bloomberg Philanthropies, and the Cancer Grand Challenges OPTIMISTICC team grant [A27140] funded by Cancer Research UK (C.L.S.).

Fig 2A and Fig S8A were created with https://biorender.com.

## Author contributions

Conceptualization: J.Q., J.L.D., C.L.S.; Methodology: J.Q., J.W., J.R.W, J.L.D, C.L.S.; Software: J.R.W.; Formal Analysis: J.Q., J.R.W., J.L.D.; Validation: J.Q., J.R.W.; Investigation: J.Q., Z.C., H.M., A.N., T.S., J.F., M.M., T.I., J.V., A.R., M.F.L., J.W., J.R.W., J.L.D.; Resources: J.Q., Z.C., J.V., J.L.D., C.L.S.; Data Curation: J.W., J.R.W., J.L.D., C.L.S.; Writing – Original Draft: J.Q.; Writing – Review & Editing: all authors; Visualization: J.Q., Z.C., J.R.W., J.L.D.; Supervision: J.L.D., C.L.S.; Project Administration: J.W., C.L.S.; Funding Acquisition: J.Q., J.L.D., C.L.S.

## Methods

### Study Participants

This study was approved by the Johns Hopkins Institutional Review Board and the University of Malaya Medical Centre (UMMC, Kuala Lumpur, Malaysia) Medical Ethics Committee. Written informed consents (provided in Malay and English) were obtained from all participants. All samples were obtained in accordance with the Health Insurance Portability and Accountability Act. Individuals who had received pre-operative radiation, chemotherapy or had a personal history of CRC were excluded. All individuals in the study underwent a standard mechanical bowel preparation. Standard pre-operative intravenous, but not oral, antibiotics were administered in all surgical cases. We collected data on the following variables: age, sex, ethnicity, BMI, smoking history, alcohol consumption, tumor location, tumor pathology, tumor grade, and tumor stage. Individuals under age 50 were defined as having early-onset CRC (EO-CRC), whereas individuals aged 50 and over were defined as having late-onset CRC (LO-CRC). Study data were collected and managed using REDCap electronic data capture tools hosted at Johns Hopkins University.

### Sample collection

Excess colon tumor and paired normal tissues were collected for analysis from individuals undergoing surgery at UMMC. Surgical tissues were fixed in methacarn or flash frozen for later analysis.

### Fluorescence *in situ* hybridization (FISH) analysis of biofilms

FISH was performed on methacarn-fixed, paraffin-embedded tissue sections as previously described ^10^. Briefly, sequential sections were stained with the Eub338 universal bacterial probe for the presence of biofilms. Serial sections of samples confirmed to have biofilms were stained with specific bacterial probes for *Fusobacterium*, *Bacteroidetes*, *Betaproteobacteria*, *Gammaproteobacteria*, and *Lachnospiraceae* (see Oligonucleotide Table). Quantification of bacteria, lambda scanning, and linear unmixing were performed on a Zeiss 780 laser scanning confocal microscope as previously described ^10^. A biofilm was defined as a dense aggregation of bacteria, with at least 20 bacteria within 1 μm of the epithelium, covering an expanse of at least 200 μm of the CEC layer, in at least 1 of 3 screened areas.

### 16S rRNA gene Illumina library generation and sequencing

16S rRNA amplicon sequencing of cohorts MAL1, MAL2 and MAL3 were performed at different facilities with different methodologies for the three sets of samples. For each cohort, the V3-V4 hypervariable region of the 16S rRNA gene was amplified and sequenced.

For the MAL1 cohort, samples were lysed by suspending in 700 μL PBS and incubating with lysozyme (5 μL of 10 mg/mL stock), mutanolysin (15 μL of 1 mg/mL stock), and lysostaphin (5 μL of 1 mg/mL stock) in lysing matrix tubes at 37°C. After 30 min, 10 μL Proteinase K and 50 μL 10% SDS was added, and samples were vortexed and then incubated at 55°C for 45 min. Mechanical lysis was performed in a FastPrep-24 5G instrument at 6.0 m/s for 40 sec, then centrifuged at 10,000 *g* for 3 min. The ZR Fecal DNA MiniPrep kit (Zymo Research) was then used for DNA extraction. The V3-V4 region was amplified using primers 319 F (5′-ACTCCTACGGGAGGCAGCAG-3′) and 806 R (5′-GGACTACHVGGGTWTCTAAT-3′) containing a linker sequence required for Illumina MiSeq 300 bp paired-end sequencing and a 12-bp heterogeneity-spacer index sequence. Sequencing was performed on the Illumina MiSeq (Illumina, San Diego, CA) according to the manufacturer’s protocol.

For the MAL2 cohort, samples were lysed and DNA extracted using the MasterPure DNA Purification Kit (Epicentre/Illumina) according to the manufacturer’s instruction. Primers used for amplification were S-D-Bact-0341-b-S-17 forward (5′-NNNNCCTACGGGNGGCWGCAG-3′) and S-D-Bact-0785-a-A-21 reverse (5’-GACTACHVGGGTATCTAATCC-3’), designed to include Illumina-compatible adaptors. Sequencing was performed on the MiSeq System (Illumina, San Diego, CA, USA) at the Monash University Malaysia Genomics Facility using the MiSeq 500-cycle reagent kit V2 on standard flow cell.

For the MAL3 cohort, DNA was extracted using the ZR Quick-DNA Fecal/Soil Microbe 96 Kit or Miniprep Kit (Zymo Research) with the following modification: 1.0mm Zirconia-Silicate beads (BioSpec) were added to the 0.1/0.5mm beads provided in the kit for improved tissue homogenization. Samples were homogenized and lysed using the Mini-beadbeater 96 (BioSpec) at maximum speed for 60 sec x 3. Lysate was then processed according to manufacturer instructions. Sequencing was performed at Novogene with DNA amplified with primers 341F (5’-CCTAYGGGRBGCASCAG-3’) and 806R (5’-GGACTACNNGGGTATCTAAT-3’), and sequencing performed on the Illumina NovaSeq 6000 platform.

### Analysis of all 16S rRNA amplicon sequence data sets

Paired-end 16S rRNA amplicon sequences were trimmed for quality and length using Trimmomatic, merged using FLASH (Fast Length Adjustment of Short reads) ^36^, and quality screened using QIIME ^37^. Spurious hits to the PhiX control genome were identified using BLASTN (Basic Local Alignment Search Tool) and removed. Passing sequences were trimmed of 16S rRNA primers, evaluated for PCR chimeras and filtered for host-associated contaminant using Bowtie2^38^. Chloroplast contaminants were identified and filtered using the RDP (Ribosomal Database Project) classifier^39^ with a confidence threshold of 50%. High-quality 16S rRNA amplicon sequences were assigned to a high-resolution taxonomic lineage using Resphera Insight, which uses a hybrid global-local alignment strategy to a manually curated 16S rRNA database with 11,000 unique species ^40–42^. This approach attempts to achieve species-level resolution when possible; however, when a confident single species assignment is not feasible, the method minimizes false positives by providing “ambiguous assignments” i.e. a list of candidate species reflecting the ambiguity. Taxonomic assignments are presented as unambiguous assignments by Resphera Insight unless otherwise indicated. Normalization of observations were performed by rarefaction followed by alpha and beta diversity characterization. Taxonomic abundances were converted from counts to percentages. Statistical associations with beta-diversity utilized PERMANOVA. Functional inference of microbial gene content was evaluated using PICRUSt2^43^. Taxonomic profiles for samples reflecting replicates were compared pairwise for quantitative concordance of detected genera and species and using Pearson and Spearman and correlation. Classification of *Fusobacterium nucleatum* C1 or C2 subgroups was performed for individual 16S rRNA amplicon sequences, using BLASTN (ungapped alignment; word size 124; minimum 99% identity) against a reference database of C1 and C2 amplicon sequence variant representatives ^12^. Candidate sequences with alignments exclusively to C1 or C2 were labeled accordingly.

### Subspecies PCRs

DNA was extracted from 3 mm tumor punch biopsies using the ZR Quick-DNA Fecal/Soil Miniprep Kit (Zymo Research) with the following modification: 1.0 mm Zirconia-Silicate beads (BioSpec) were added to the 0.1/0.5 mm beads provided in the kit for improved tissue homogenization. Samples were homogenized and lysed using the Mini-beadbeater 96 (BioSpec) at maximum speed for 60 sec x 3. Lysate was then processed according to manufacturer instructions. DNA concentration was measured on a nanodrop, and ranged from 30.4 to 260.5 ng/μl. DNA was screened for *F. nucleatum* using primers specific for *nusG* (forward, 5’-CAACCATTACTTTAACTCTACCATGTTCA-3’; reverse, 5’-GTTGACTTTACAGAAGGAGATTATGTAAAAATC-3’)^3^ at a primer concentration of 0.9 μM using a PCR program of 98°C x 60 sec; 35 cycles of 98°C x 30 sec, 58°C x 60 sec, 72°C x20 sec; and 72°C x 7 min. DNA was then screened using primers specific for each *F. nucleatum* subspecies (See Oligonucleotide Table) ^16^ at a primer concentration of 0.5 μM, using a PCR program of 98°C x 60 sec; 30 cycles of 98°C x 30 sec, 56°C x 15 sec, 72°C x20 sec; and 72°C x 7 min.

### Selective Culture

Three mm punches were taken from flash-frozen tumor biopsy tissues and homogenized with a sterile pestle in 50 μl brain heart infusion (BHI) media supplemented with hemin (10 μg/mL) and vitamin K (5 μg/mL). Total volume was brought up to 500 μl of BHI and vortexed. 10 μl of the homogenate were spread onto *Fusobacterium* selective agar (FSA; Anaerobe Systems) and 10 μl were subcultured in 3 mL of BHI. Each was incubated for 48-72 hr under anaerobic conditions (75% N2, 5% H2, 20% CO2) in an anaerobic chamber. From the homogenate culture in BHI media, 10 μl were then streaked onto FSA plates and incubated as above. From these two FSA plates (1 directly plated, 1 subcultured first in BHI), unique colonies were picked and spread onto new FSA plates for isolation (minimum 6 colonies/tumor) and incubated. After 48-72 hr, colonies were scooped up and swirled into 50 μl sterile water, which was boiled to release bacterial DNA. This crude DNA prep was used as a template for screening by quantitative real-time PCR (qRT-PCR) with *Fusobacterium* genus-specific 16S primers (forward, GGATTTATTGGGCGTAAAG; reverse, GGCATTCCTACAAATATCTACGAA)^44^ and probe (5’VIC-TGC AGG GCT CAA CTC TGT ATT GCG - 3’BHQ1), at a concentration of 0.2 μM, and with the following program: 50°C x 2 min; 95°C x 10 min 40 cycles of 95°C x 15 sec, 58°C x 60 sec. Colonies were considered positive if the CT value was <30. Each positive colony was subcultured into BHI media x 48 hr, and genomic DNA was extracted from pelleted bacteria using the ZR Quick-DNA Fecal/Soil Miniprep Kit (Zymo Research). Species and subspecies were confirmed by Sanger sequencing of the 16S rRNA gene (Azenta Life Sciences).

### Quantification and statistical analysis

To compare categorical data between defined groups (e.g. biofilm status by tumor location), data were analyzed using a two-tailed Mann Whitney non-parametric t-test. Comparison of biofilm subtype across tumor stages was analyzed by a nonparametric Kruskal-Wallis one-way ANOVA. Prior to downstream statistical comparisons, 16S rRNA amplicon profiles within each dataset were subsampled to an even level of coverage (9500 reads). For comparisons of relative taxonomic abundances between matched tumor-normal pairs, data were analyzed by a two-tailed non-parametric Wilcoxon signed-rank sum test. To identify associations of oral taxa abundances with *F. nucleatum* abundance, we performed a non-parametric Spearman correlation. Differences in PICRUSt2 pathway content was analyzed by Mann Whitney Test or by Kruskal-Wallis test with Dunn’s multiple comparisons, as indicated in the figure legend. For all analyses, differences with a P value of <0.05 were considered significant.

### Data and code availability

Raw sequences from MAL1 and MAL2 were previously deposited in the NCBI SRA repository under BioProject accession nos. PRJNA325649 and PRJNA325650, respectively. Raw sequences from MAL3 were deposited in the NCBI SRA repository under BioProject accession no. PRJNA1195962. Primary sequencing data and open source R code used for meta-analyses are available from the authors upon request.

### Oligonucleotide Table

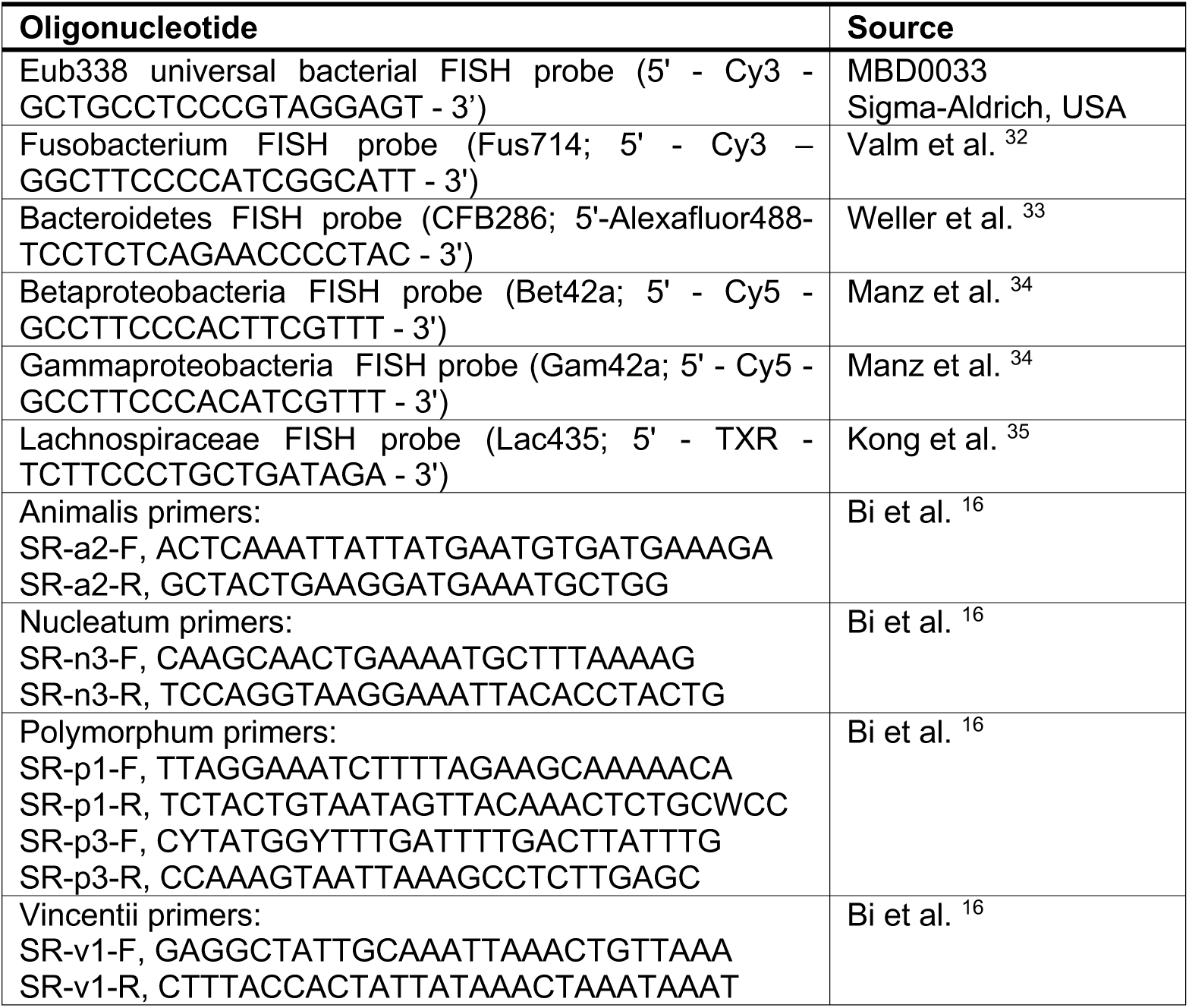

