## Supplemental Figures for "*Fusobacterium nucleatum* is enriched in invasive biofilms in colorectal cancer"

### Figure S1

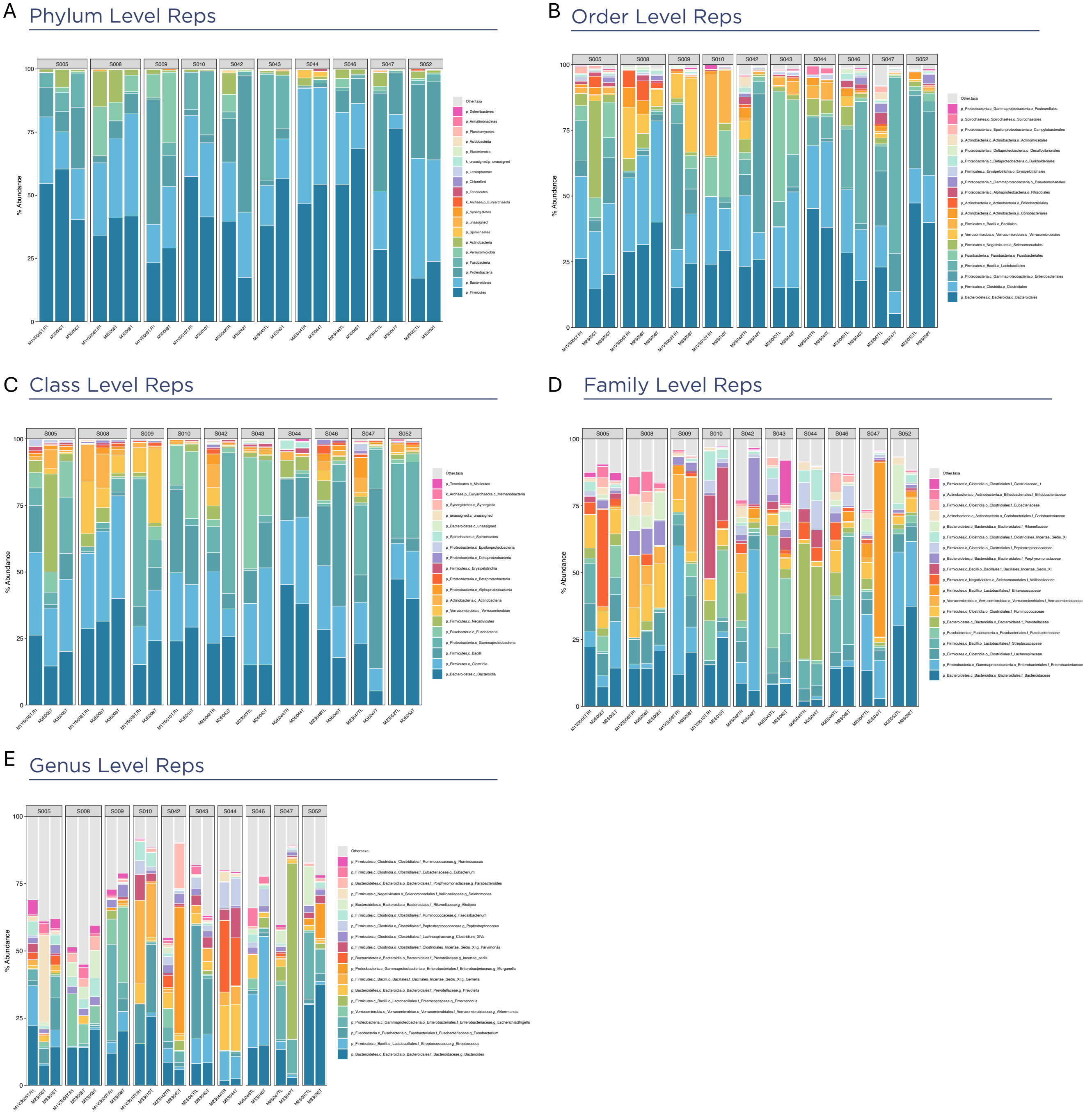
A
Phylum Level Reps
B
Order Level Reps
C
Class Level Reps
D
Family Level Reps
E
Genus Level Reps
% Abundance

**Figure S1. Relative abundance of sequencing cohort replicates.** Relative abundance by 16S rRNA amplicon sequencing of 10 tumors sequenced in replicate between cohorts MAL1, MAL2, and/or MAL3 at the A) phylum, B) order, C) Class, D) family, and E) genus level.

Figure S2

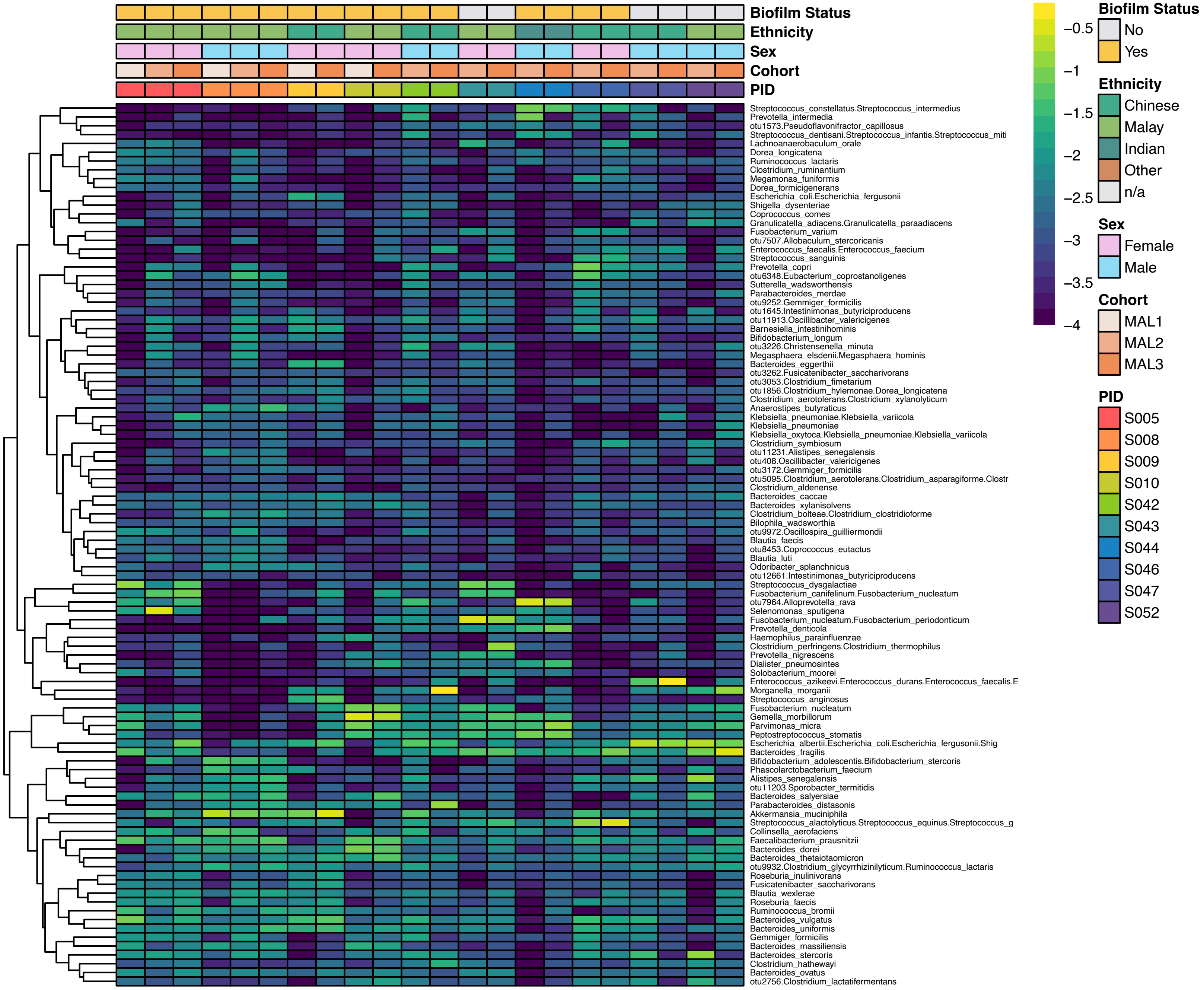

Biofilm Status
Ethnicity
Sex
Cohort
PID
Streptococcus_constellatus.Streptococcus_intermedius
Prevotella_intermedia
otu1573.Pseudoflavonifractor_capillosus
Streptococcus_dentisani.Streptococcus_infantis.Streptococcus_miti
Lachnoanaerobaculum_orale
Dorea_longicatena
Ruminococcus_lactaris
Clostridium_ruminantium
Megamonas_funiformis
Dorea_formicigenerans
Escherichia_coli.Escherichia_fergusonii
Shigella_dysenteriae
Coprococcus_comes
Granulicatella_adiacens.Granulicatella_paraadiacens
Fusobacterium_varium
otu7507.Allobaculum_stercoricanis
Enterococcus_faecalis.Enterococcus_faecium
Streptococcus_sanguinis
Prevotella_copri
otu6348.Eubacterium_coprostanoligenes
Sutterella_wadsworthensis
Parabacteroides_merdae
otu9252.Gemmiger_formicilis
otu1645.Intestinimonas_butyriciproducens
otu11913.Oscillibacter_valericigenes
Barnesiella_intestinihominis
Bifidobacterium_longum
otu3226.Christensenella_minuta
Megasphaera_elsdenii.Megasphaera_hominis
Bacteroides_eggerthii
otu3262.Fusicatenibacter_saccharivorans
otu3053.Clostridium_fimetarium
otu1856.Clostridium_hylemonae.Dorea_longicatena
Clostridium_aerotolerans.Clostridium_xylanolyticum
Anaerostipes_butyraticus
Klebsiella_pneumoniae.Klebsiella_variicola
Klebsiella_pneumoniae
Klebsiella_oxytoca.Klebsiella_pneumoniae.Klebsiella_variicola
Clostridium_symbiosum
otu11231.Alistipes_senegalensis
otu408.Oscillibacter_valericigenes
otu3172.Gemmiger_formicilis
otu5095.Clostridium_aerotolerans.Clostridium_asparagiforme.Clostr
Clostridium_aldenense
Bacteroides_caccae
Bacteroides_xylanisolvens
Clostridium_bolteae.Clostridium_clostridioforme
Bilophila_wadsworthia
otu9972.Oscillospira_guilliermondii
Blautia_faecis
otu8453.Coprococcus_eutactus
Blautia_luti
Odoribacter_splanchnicus
otu12661.Intestinimonas_butyriciproducens
Streptococcus_dysgalactiae
Fusobacterium_canifelinum.Fusobacterium_nucleatum
otu7964.Alloprevotella_rava
Selenomonas_sputigena
Fusobacterium_nucleatum.Fusobacterium_periodonticum
Prevotella_denticola
Haemophilus_parainfluenzae
Clostridium_perfringens.Clostridium_thermophilus
Prevotella_nigrescens
Dialister_pneumosintes
Solobacterium_moorei
Enterococcus_azikeevi.Enterococcus_durans.Enterococcus_faecalis.E
Morganella_morganii
Streptococcus_anginosus
Fusobacterium_nucleatum
Gemella_morbillorum
Parvimonas_micra
Peptostreptococcus_stomatis
Escherichia_albertii.Escherichia_coli.Escherichia_fergusonii.Shig
Bacteroides_fragilis
Bifidobacterium_adolescentis.Bifidobacterium_stercoris
Phascolarctobacterium_faecium
Alistipes_senegalensis
otu11203.Sporobacter_termitidis
Bacteroides_salyersiae
Parabacteroides_distasonis
Akkermansia_muciniphila
Streptococcus_alactolyticus.Streptococcus_equinus.Streptococcus_g
Collinsella_aerofaciens
Faecalibacterium_prausnitzii
Bacteroides_dorei
Bacteroides_thetaiotaomicron
otu9932.Clostridium_glycyrrhizinilyticum.Ruminococcus_lactaris
Roseburia_inulinivorans
Fusicatenibacter_saccharivorans
Blautia_wexlerae
Roseburia_faecis
Ruminococcus_bromii
Bacteroides_vulgatus
Bacteroides_uniformis
Gemmiger_formicilis
Bacteroides_massiliensis
Bacteroides_stercoris
Clostridium_hathewayi
Bacteroides_ovatus
otu2756.Clostridium_lactatifermentans
−0.5
−1
−1.5
−2
−2.5
−3
−3.5
−4
Biofilm Status
No
Yes
Ethnicity
Chinese
Malay
Indian
Other
n/a
Sex
Female
Male
Cohort
MAL1
MAL2
MAL3
PID
S005
S008
S009
S010
S042
S043
S044
S046
S047
S052

**Figure S2. Species abundance of sequencing cohort replicates.** Heatmap depicting relative abundance by 16S rRNA amplicon sequencing of 10 tumors sequenced in replicate between cohorts MAL1, MAL2, and/or MAL3 at the species level, with metadata color-coded as depicted in the legend for biofilm status, ethnicity, sex, sequencing cohort, and patient identification number (PID).

Figure S3

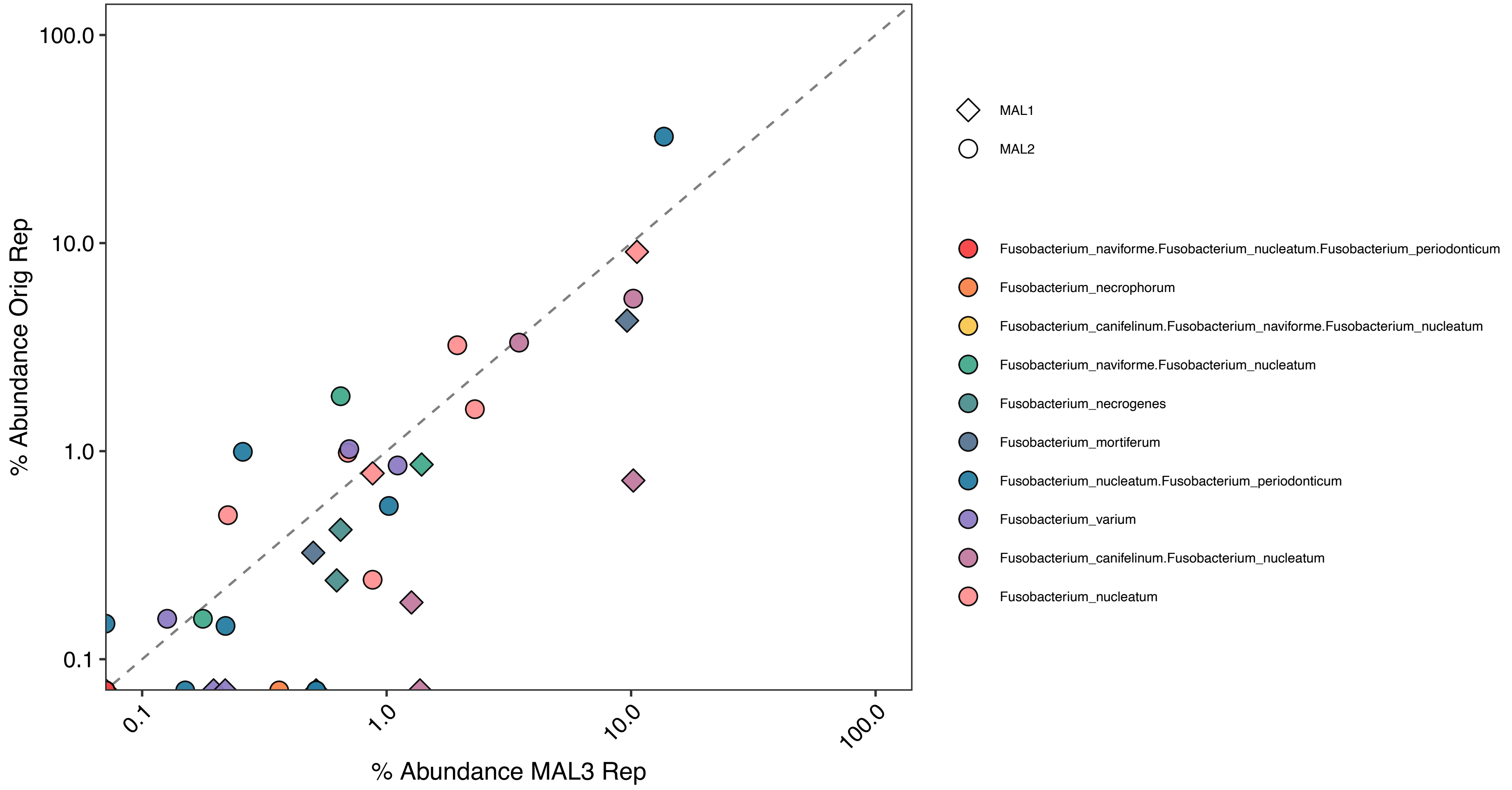
100.0
10.0
1.0
0.1
% Abundance Orig Rep
0.1
1.0
10.0
100.0
% Abundance MAL3 Rep
MAL1
MAL2
Fusobacterium_naviforme.Fusobacterium_nucleatum.Fusobacterium_periodonticum
Fusobacterium_necrophorum
Fusobacterium_canifelinum.Fusobacterium_naviforme.Fusobacterium_nucleatum
Fusobacterium_naviforme.Fusobacterium_nucleatum
Fusobacterium_necrogenes
Fusobacterium_mortiferum
Fusobacterium_nucleatum.Fusobacterium_periodonticum
Fusobacterium_varium
Fusobacterium_canifelinum.Fusobacterium_nucleatum
Fusobacterium_nucleatum

**Figure S3. Relative *Fusobacterium species* abundance of sequencing cohort replicates.** Relative abundance of *Fusobacterium* species by 16S rRNA amplicon sequencing of 10 tumors sequenced in replicate between cohorts MAL1, MAL2, and/or MAL3 , depicted as percent abundance of the original replicate in MAL1 or MAL2 graphed against percent abundance of the MAL3 replicate. Ambiguous assignments depicted in the legend indicate inability to resolve two or more species (e.g. "Fusobacterium_nucleatum.Fusobacterium_peridonticum" indicates that sequences could not distinguish between *F. nucleatum* and *F. peridonticum*).

Figure S4

A

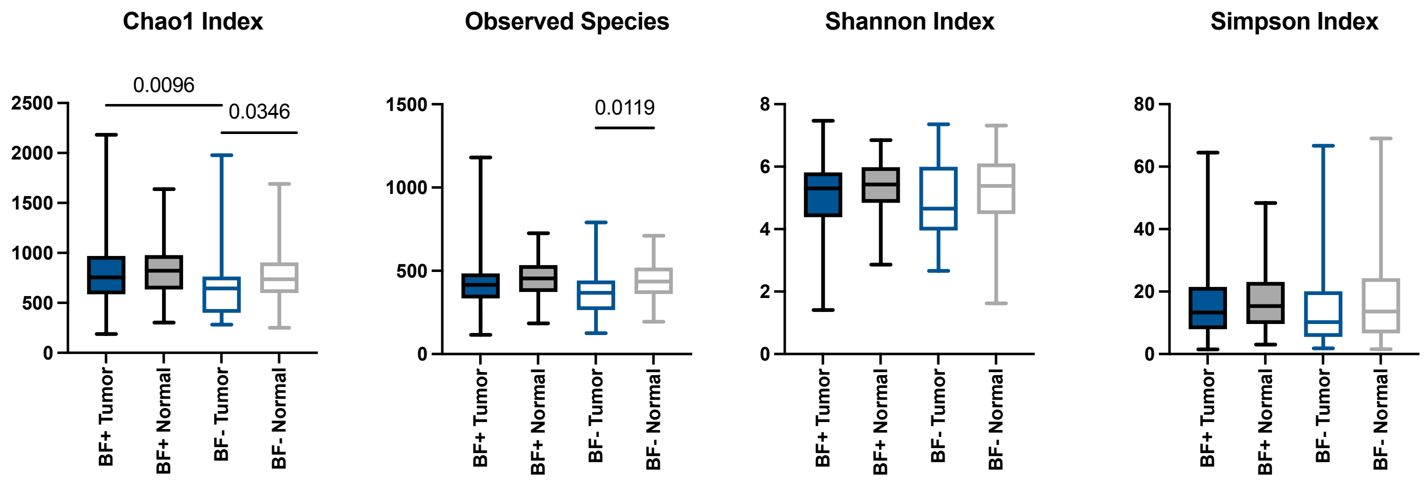
Chao1 Index
0.0096
0.0346
2500
2000
1500
1000
500
0
BF+ Tumor
BF+ Normal
BF- Tumor
BF- Normal
Observed Species
0.0119
1500
1000
500
0
Shannon Index
8
6
4
2
0
Simpson Index
80
60
40
20
0

B

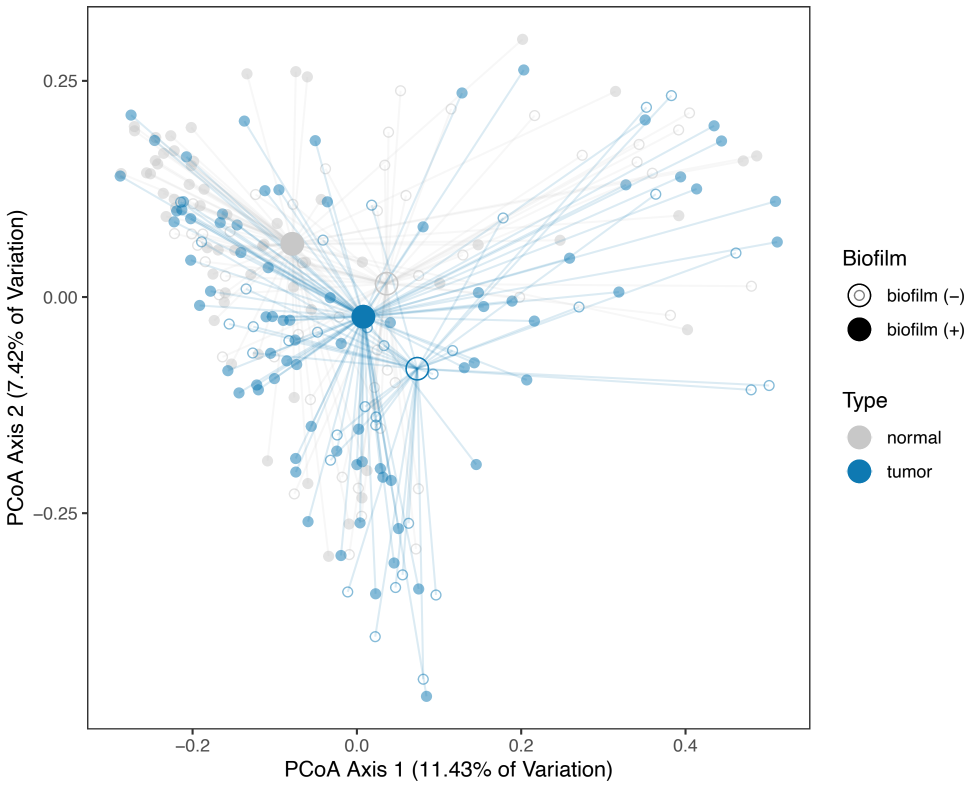
PCoA Axis 1 (11.43% of Variation)
PCoA Axis 2 (7.42% of Variation)
0.25
0.00
−0.25
−0.2
0.0
0.2
0.4
Biofilm
biofilm (−)
biofilm (+)
Type
normal
tumor

**Figure S4. Alpha and beta diversity of tumor and normal tissue by biofilm status.** A) Measures of alpha diversity for tumor and normal samples with or without biofilms. BF+ indicates biofilm-positive and BF- indicates biofilm-negative. Box and whisker plots depict min to max. Analyzed by two-tailed Mann Whitney test, with $p<0.05$ considered significant. B) Principal coordinate analysis plots (PCoA) of Bray Curtis dissimilarity depicted as centroids. Analyzed by PERMANOVA, with $p<0.05$ considered significant.

Figure S5

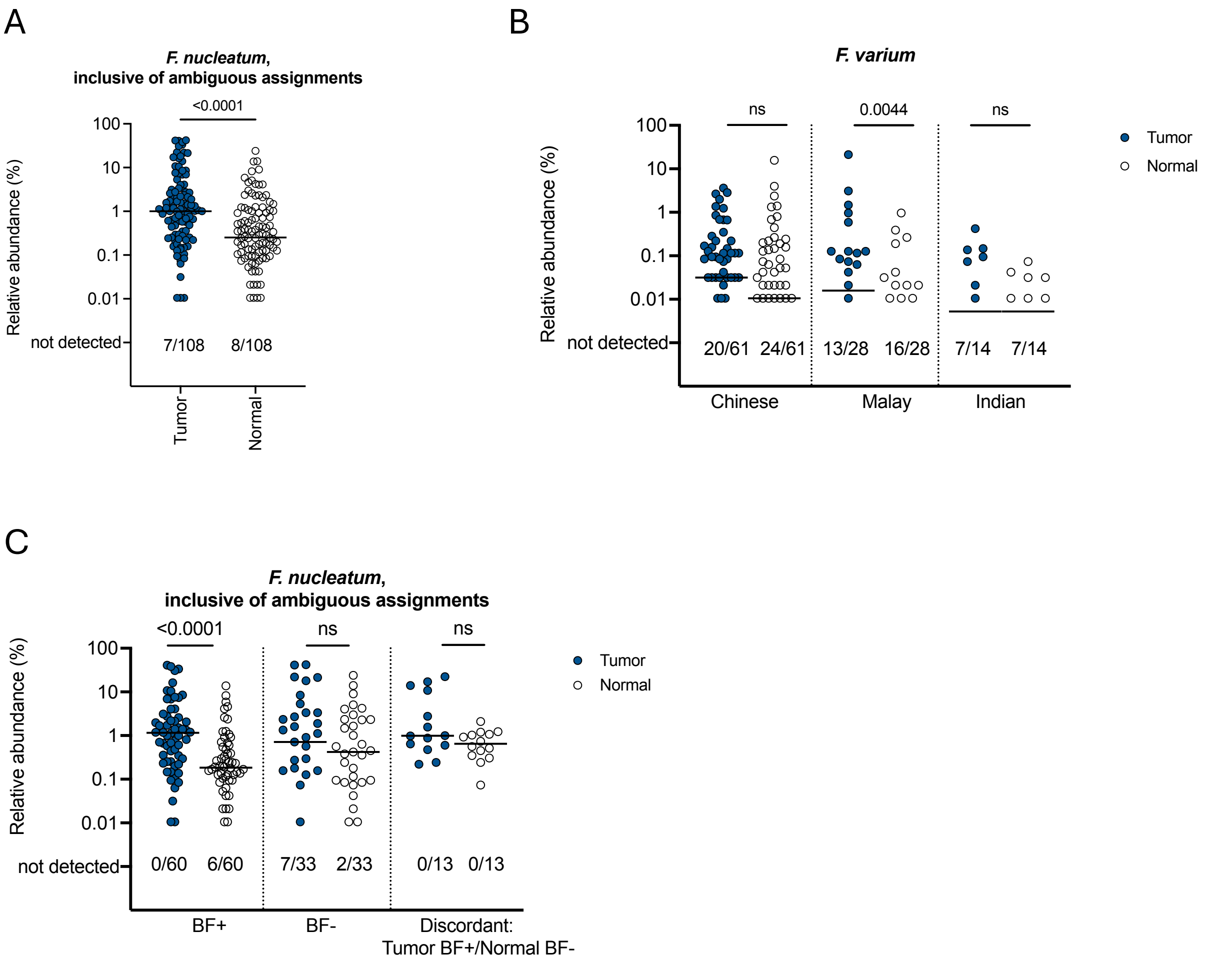
A
F. nucleatum,
inclusive of ambiguous assignments
<0.0001
Relative abundance (%)
100
10
1
0.1
0.01
not detected
7/108
8/108
Tumor
Normal
B
F. varium
ns
0.0044
ns
Relative abundance (%)
100
10
1
0.1
0.01
not detected
20/61
24/61
13/28
16/28
7/14
7/14
Chinese
Malay
Indian
Tumor
Normal
C
F. nucleatum,
inclusive of ambiguous assignments
<0.0001
ns
ns
Relative abundance (%)
100
10
1
0.1
0.01
not detected
0/60
6/60
7/33
2/33
0/13
0/13
BF+
BF-
Discordant:
Tumor BF+/Normal BF-
Tumor
Normal

**Figure S5. Additional details on *Fusobacterium nucleatum* and *Fusobacterium varium* abundance.** A, C) Relative abundance by 16S rRNA amplicon sequencing of each depicted *Fusobacterium nucleatum* as a sum of confident and ambiguous species assignments in pairs of tumor (blue circle) and normal (clear circle) tissues, stratified by biofilm (BF) status in panel C. B) Relative abundance of *Fusobacterium varium* abundance stratified by ethnicity in pairs of tumor (blue circle) and normal (clear circle) tissues. Bars indicate median. Numbers below each graph depict the number of samples out of the total in which each species was not detected by sequencing. Analyzed by two-tailed Wilcoxon matched-pairs signed rank test, with $p<0.05$ considered significant.

### Figure S6

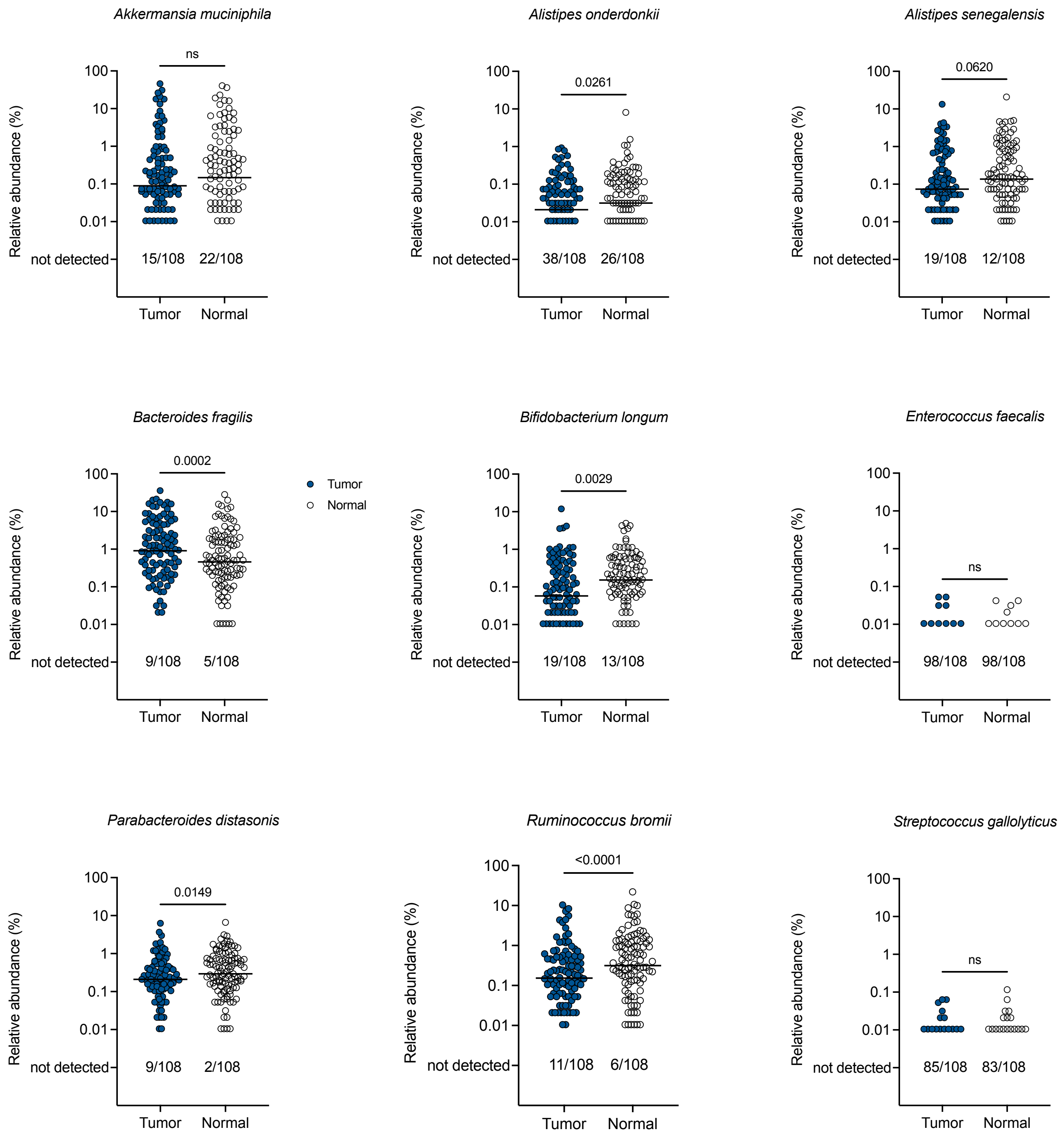
Akkermansia muciniphila
ns
15/108
22/108
Alistipes onderdonkii
0.0261
38/108
26/108
Alistipes senegalensis
0.0620
19/108
12/108
Bacteroides fragilis
0.0002
9/108
5/108
Tumor
Normal
Bifidobacterium longum
0.0029
19/108
13/108
Enterococcus faecalis
ns
98/108
98/108
Parabacteroides distasonis
0.0149
9/108
2/108
Ruminococcus bromii
<0.0001
11/108
6/108
Streptococcus gallolyticus
ns
85/108
83/108
Relative abundance (%)
100
10
1
0.1
0.01
not detected
Tumor
Normal

**Figure S6**. **Abundance of other CRC-associated species.** Relative abundance by 16S rRNA amplicon sequencing of each depicted species in pairs of tumor (blue circle) and normal (clear circle) tissues. Bars indicate median. Numbers below each graph depict the number of samples out of the total in which each species was not detected by sequencing. Analyzed by two-tailed Wilcoxon matched-pairs signed rank test, with $p<0.05$ considered significant.

Figure S7

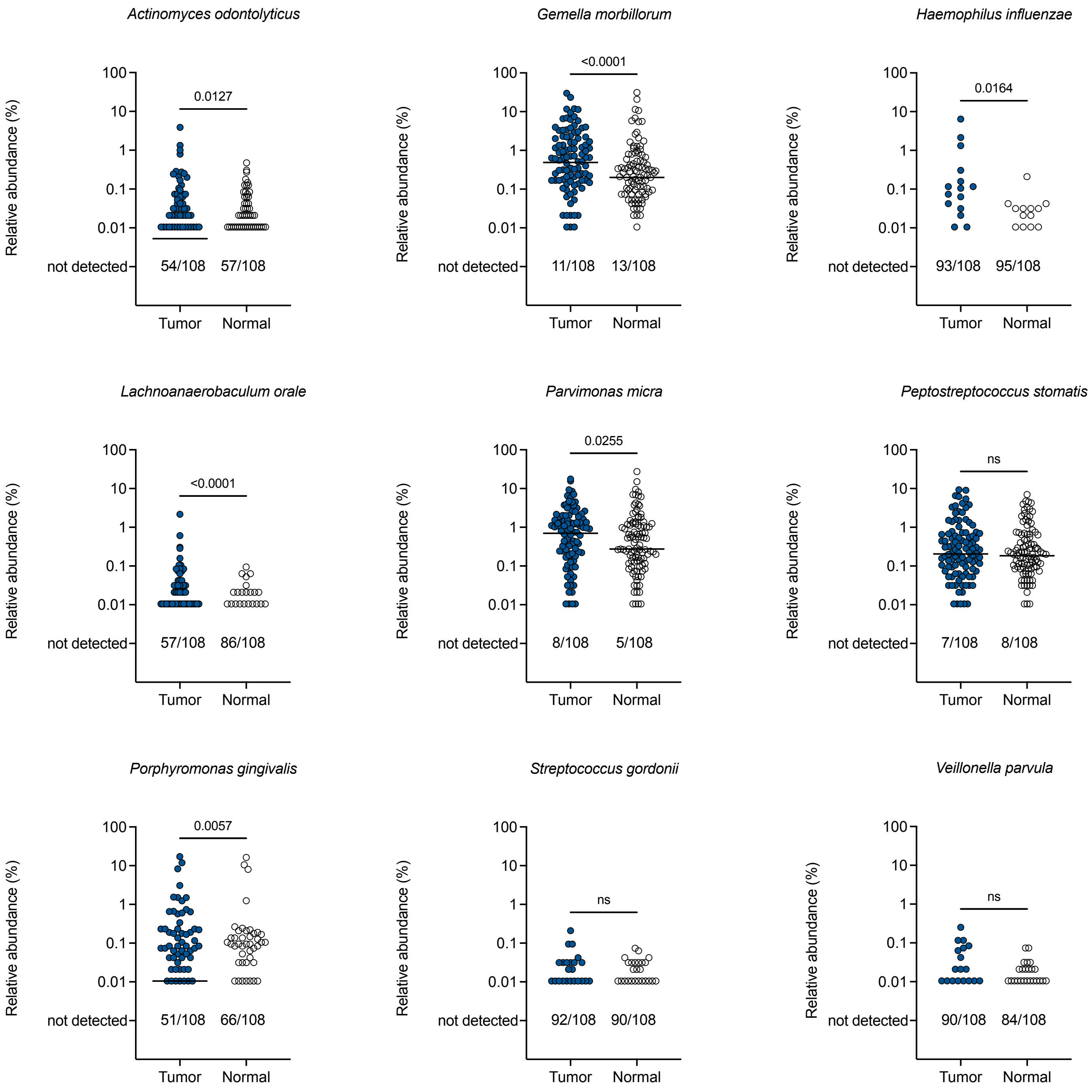

Actinomyces odontolyticus
Relative abundance (%)
100
10
1
0.1
0.01
not detected
0.0127
54/108
57/108
Tumor
Normal
Gemella morbillorum
<0.0001
11/108
13/108
Haemophilus influenzae
0.0164
93/108
95/108
Lachnoanaerobaculum orale
<0.0001
57/108
86/108
Parvimonas micra
0.0255
8/108
5/108
Peptostreptococcus stomatis
ns
7/108
8/108
Porphyromonas gingivalis
0.0057
51/108
66/108
Streptococcus gordonii
ns
92/108
90/108
Veillonella parvula
ns
90/108
84/108

**Figure S7. Abundance of oral biofilm organisms in CRC tumors/normal pairs.** Relative abundance by 16S rRNA amplicon sequencing of each depicted species in pairs of tumor (blue circle) and normal (clear circle) tissues. Bars indicate median. Numbers below each graph depict the number of samples out of the total in which each species was not detected by sequencing. Analyzed by two-tailed Wilcoxon matched-pairs signed rank test, with $p<0.05$ considered significant.

Figure S8

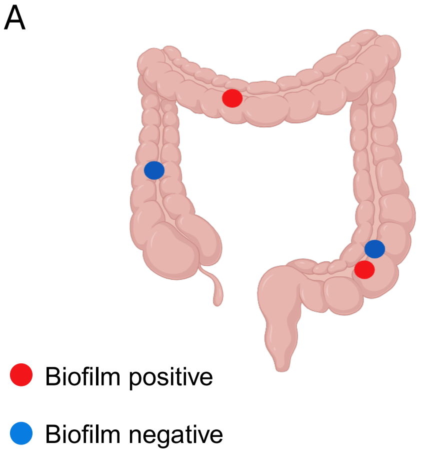

A
Biofilm positive
Biofilm negative


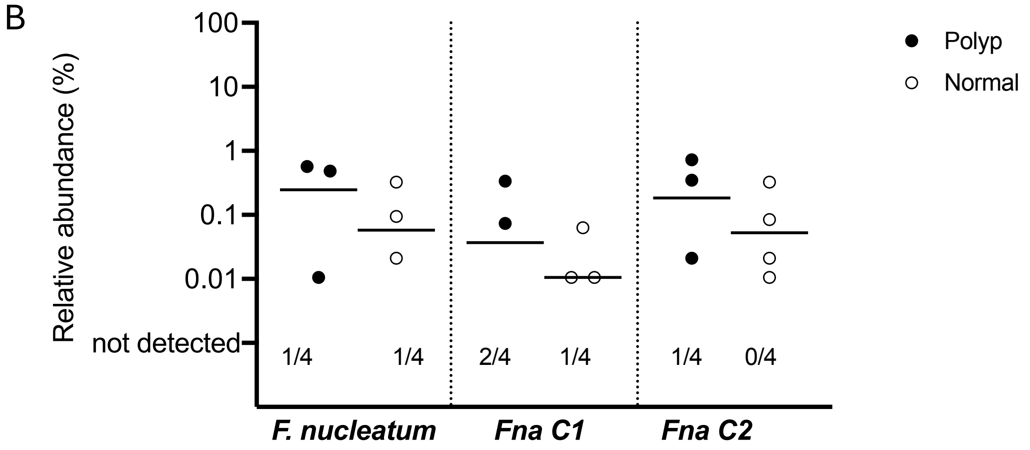

B
Relative abundance (%)
100
10
1
0.1
0.01
not detected
1/4
1/4
2/4
1/4
1/4
0/4
F. nucleatum
Fna C1
Fna C2
Polyp
Normal

**Figure S8. Adenomas in the cohort.** A) Diagram of adenoma (polyp) location in the colon and biofilm status (red circle = biofilm-positive; blue circle = biofilm-negative). B) Relative abundance by 16S rRNA amplicon sequencing of *Fusobacterium nucleatum*, *subspecies animalis* clade 1 (Fna C1) and clade 2 (Fna C2) in pairs of polyp (black circle) and normal (clear circle) tissues. Bars indicate median. Numbers below each graph depict the number of samples out of the total in which each species was not detected by sequencing.

Figure S9

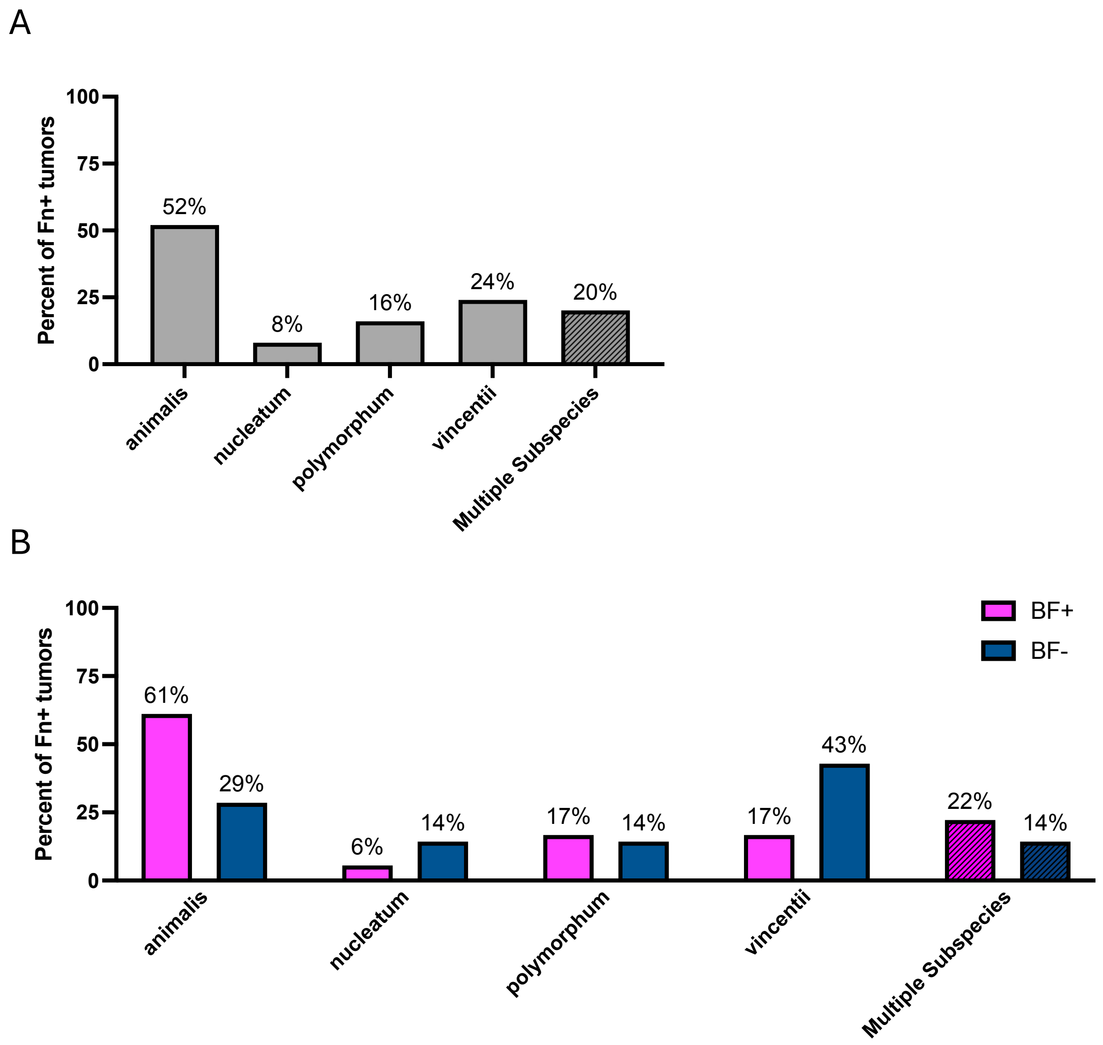
A
100
75
50
25
0
Percent of Fn+ tumors
52%
8%
16%
24%
20%
animalis
nucleatum
polymorphum
vincentii
Multiple Subspecies
B
100
75
50
25
0
Percent of Fn+ tumors
BF+
BF-
61%
29%
6%
14%
17%
14%
17%
43%
22%
14%
animalis
nucleatum
polymorphum
vincentii
Multiple Subspecies

**Figure S9. *Fusobacterium nucleatum* subspecies detected in tumors by PCR.** Tumor DNA samples positive for *F. nucleatum* (Fn+) using primers for *nusG* (n=25) were screened with subspecies-specific PCRs [16]. A) Percent of these tumors in which bands were detected for each subspecies or for multiple subspecies, and B) stratified by the biofilm (BF) status of the tumor.

Figure S10

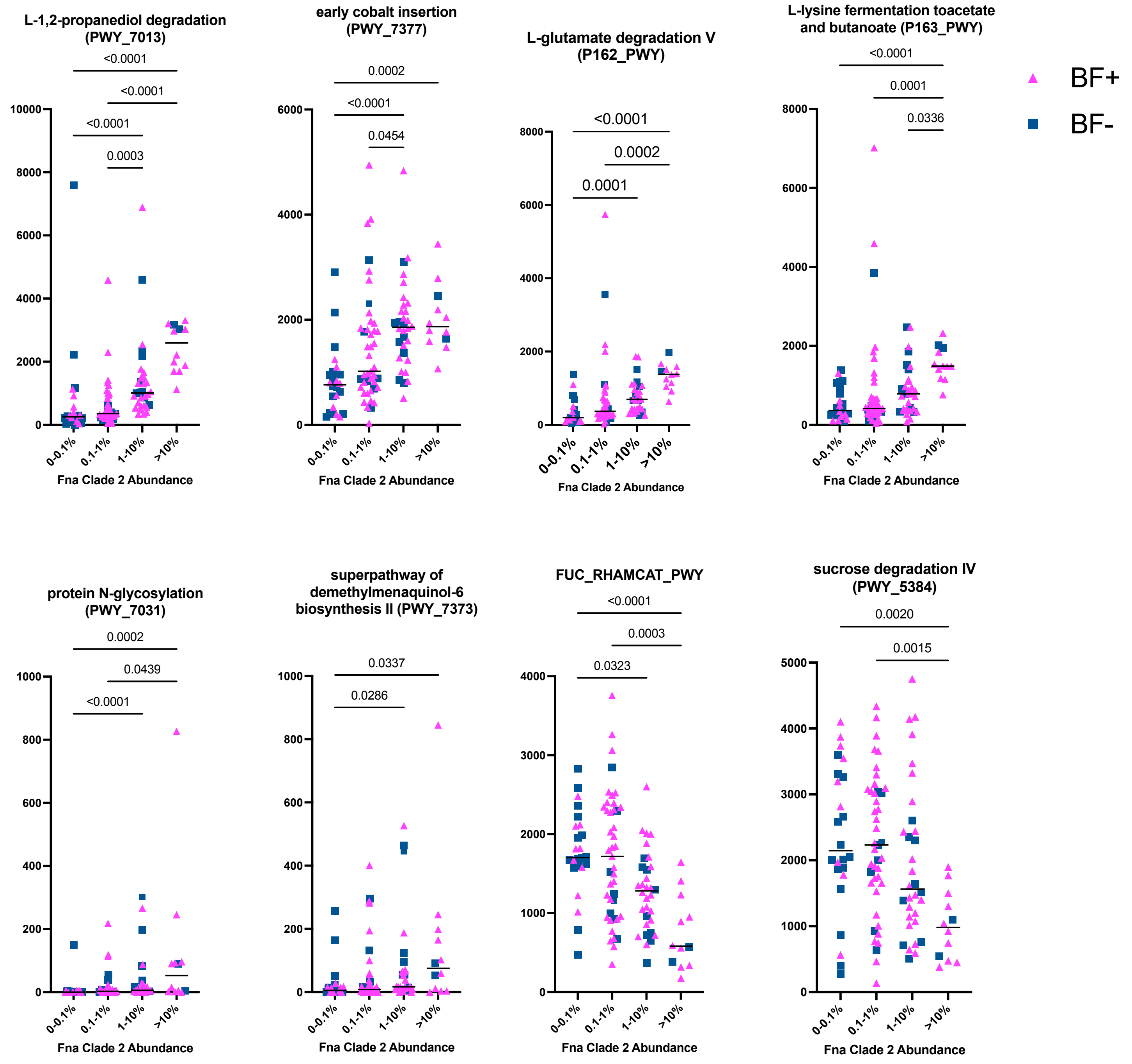

BF+
BF-
L-1,2-propanediol degradation (PWY_7013)
<0.0001
<0.0001
<0.0001
0.0003
early cobalt insertion (PWY_7377)
0.0002
<0.0001
0.0454
L-glutamate degradation V (P162_PWY)
<0.0001
0.0002
0.0001
L-lysine fermentation toacetate and butanoate (P163_PWY)
<0.0001
0.0001
0.0336
protein N-glycosylation (PWY_7031)
0.0002
0.0439
<0.0001
superpathway of demethylmenaquinol-6 biosynthesis II (PWY_7373)
0.0337
0.0286
FUC_RHAMCAT_PWY
<0.0001
0.0003
0.0323
sucrose degradation IV (PWY_5384)
0.0020
0.0015
0-0.1%
0.1-1%
1-10%
>10%
Fna Clade 2 Abundance

**Figure S10. PICRUSt2 predictions of metabolic function associated with Fna C2 abundance in CRC tumors.** Specific PICRUSt2 pathways of interest from among those depicted in Fig 6, for biofilm positive (pink triangle) and biofilm negative (blue square) tumors for each Fna C2 abundance category. Bars indicate median. Analyzed by Kruskal-Wallis test and Dunn's multiple comparisons, with $p<0.05$ considered significant.

Figure S11

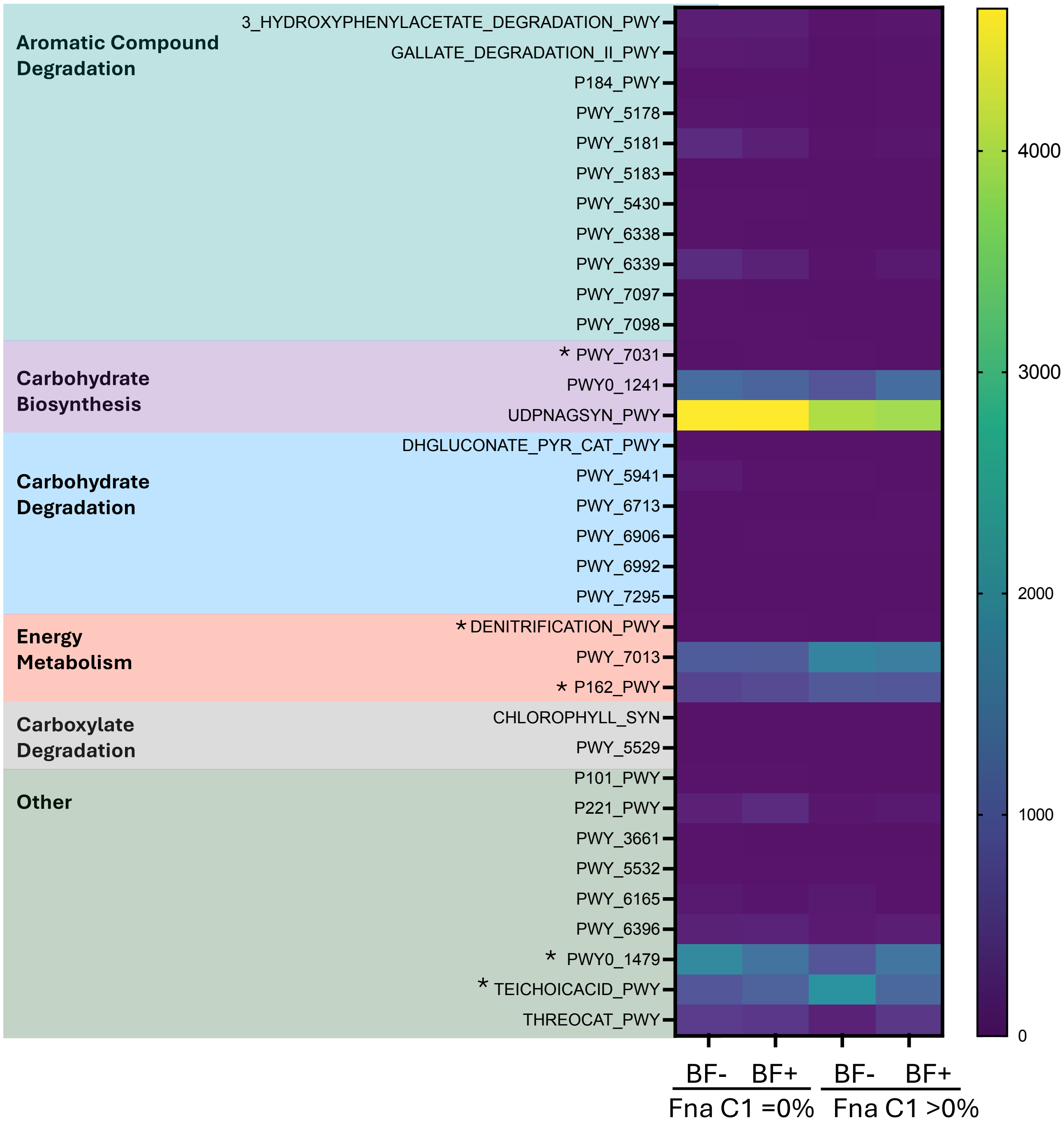
Aromatic Compound Degradation
3_HYDROXYPHENYLACETATE_DEGRADATION_PWY
GALLATE_DEGRADATION_II_PWY
P184_PWY
PWY_5178
PWY_5181
PWY_5183
PWY_5430
PWY_6338
PWY_6339
PWY_7097
PWY_7098
Carbohydrate Biosynthesis
* PWY_7031
PWY0_1241
UDPNAGSYN_PWY
Carbohydrate Degradation
DHGLUCONATE_PYR_CAT_PWY
PWY_5941
PWY_6713
PWY_6906
PWY_6992
PWY_7295
Energy Metabolism
* DENITRIFICATION_PWY
PWY_7013
* P162_PWY
Carboxylate Degradation
CHLOROPHYLL_SYN
PWY_5529
Other
P101_PWY
P221_PWY
PWY_3661
PWY_5532
PWY_6165
PWY_6396
* PWY0_1479
* TEICHOICACID_PWY
THREOCAT_PWY
4000
3000
2000
1000
0
BF- BF+ BF- BF+
Fna C1 =0%
Fna C1 >0%

**Figure S11. PICRUSt2 pathways associated with Fna C1 abundance in CRC tumors.** Heatmap of all statistically significant PICRUSt2 pathways with significant differences in gene content between absent Fna C1 (0% abundance) and present Fna C1 (>0%) in tumors (n=112), as analyzed by Mann Whitney test, with $p<0.05$ considered significant. Pathways denoted with an asterix (*) were also significantly different between Fna C2 abundance categories, as depicted in Fig 6. Data are further stratified by biofilm (BF) status within each abundance category.

**Supplementary Tables**

- Extended Data 1: 16S Sequencing results of 10 replicates
  - Table S1A: pre-processing data
  - Table S1B: phylum
  - Table S1C: class
  - Table S1D: order
  - Table S1E: family
  - Table S1F: genus
  - Table S1G: species/OTU

- Extended Data 2: 16S Sequencing results of full cohort
  - Table S2A: 16S alpha diversity
  - Table S2B: 16S phylum
  - Table S2C: 16S class
  - Table S2D: 16S order
  - Table S2E: 16S family
  - Table S2F: 16S genus
  - Table S2G: 16S species/OTU
  - Table S2H: Fna clades
  - Table S2I: PICRUSt2 pathways

- Table S3: *Fusobacterium* strains isolated by culture
